# *De Novo* Hepatic Pyrimidine Synthesis Regulates Systemic Energy Homeostasis

**DOI:** 10.1101/2025.11.18.689117

**Authors:** Ling Fu, Pradnya H. Patil, Pengfei Zhang, Ignacio Norambuena-Soto, Qiutang Xiong, Yin Wang, Chelsea Jang, Kevin Wang, Yunqian Peng, Lu Ding, Patrick Fueger, Lihua Jin, Wendong Huang, Philip L. Lorenzi, Lin Tan, Markus Kalkum, Philipp E Scherer, Zhao V. Wang, Yingfeng Deng

**Author notes:** Corresponding author: Yingfeng Deng, Ph.D. Department of Diabetes and Cancer Metabolism, Arthur Riggs Diabetes and Metabolism Research Institute, Beckman Research Institute, City of Hope National Medical Center, 1500 East Duarte Road, Duarte, California 91010, USA.

## Abstract

Fasting blood uridine is increased in obesity and type 2 diabetes (T2D), but the significance of hepatic uridine biosynthesis to the etiology of both remains elusive. We found that *de novo* pyrimidine synthesis in the liver is reduced by fasting and diet-induced obesity, while suppression of hepatic pyrimidine synthesis promotes obesity and insulin resistance. The metabolic sequalae of hepatic pyrimidine synthesis suppression, however, is not associated with altered plasma uridine concentration. Instead, it is associated with an increased hepatic glucose production and a decreased hepatic insulin clearance, two key functions of hepatocytes in regulating systemic energy homeostasis. We found that enhanced gluconeogenesis is the primary reason for increased hepatic glucose production. Moreover, uridine, which was maintained stable in the circulation by adipose tissue and the liver, preferentially shut down pyrimidine synthesis in hepatocytes but not adipocytes at blood concentrations that occur with fasting. Remarkably, uridine, at fasting levels, increases gluconeogenesis further in hepatocytes when *de novo* pyrimidine synthesis is suppressed, indicating a synergistical action of uridine and its biosynthesis pathway in promoting hepatic glucose production, a mechanism highly relevant to the pathophysiology of insulin resistance in obesity. Theologically, maintenance of blood uridine within the narrow range protects mammals from high-rate spontaneous tumorigenesis. Since obesity promotes an increase in blood uridine from adipocytes, suppressing uridine synthesis in hepatocytes becomes a critical response to lower spontaneous tumorigenesis. Pyrimidine synthesis suppression in hepatocytes, however, promotes gluconeogenesis and ultimately triggers obesity and T2D. These findings suggest a new paradigm for the etiology of metabolic deterioration in diet-induced obesity, in which perturbations in uridine promotes obesity and T2D.

## Introduction

Uridine is an uracil conjugated nucleoside and plays multifaceted roles in cells [1]. Serving as a building block for RNA and DNA biosynthesis[2], uridine is critical for cell proliferation and growth. Uridine also promotes cell growth by facilitating synthesis of sugar nucleotides, the substrates necessary for both protein and lipid glycosylation [3, 4]. Moreover, uridine modulates the insulin sensitivity in cells through the hexosamine biosynthesis pathway [5–7]. Multiple studies reported that blood uridine levels are elevated in obesity and type 2 diabetes (T2D) [8–10], however, the driving force of the elevation and its impact on glycemic control remain elusive.

In mammals, blood uridine levels are under stringent regulation in mammals [11, 12]. In Upp1 KO mice, a defect in uridine catabolism leads to excess uridine, DNA damage, and spontaneous tumorigenesis, highlighting the detrimental effect of high blood uridine [13]. We reported that the clearance of blood uridine in mice is linked to feeding as food stimulates bile excretion and therefore promotes bile-mediated uridine clearance [14]. The dynamic change in blood uridine during fasting-feeding is also found in humans [14, 15], implying that uridine homeostasis is tied to energy metabolism in mammals. Our group and others found that blood uridine levels are elevated in obese mice that are deficient in leptin signaling, a key hormonal pathway that regulates energy balance [10, 14], suggesting that energy imbalance can also lead to dysregulation of uridine homeostasis. However, the mechanistic link is unclear.

The liver has been thought responsible for blood uridine supply in mammals [16, 17]. Recently we demonstrated that adipocytes are also a key source of blood uridine [14, 18]. When circulating uridine levels are increased by fasting, pyrimidine synthesis is decreased in the liver, and adipocytes serve as the major source of uridine supply [14]. One question is why a synergy in uridine biosynthesis exists between the liver and adipose tissue. Since the adipocytes do not show a fasting-induced change in uridine synthesis, we wondered if turning off uridine synthesis is more critical to the liver as a fasting adaption. *De novo* pyrimidine synthesis consumes ribose-5-phosphate and aspartate. This biosynthesis process is unfavored when glucose production is demanded in hepatocytes. Meanwhile, since obesity is associated with increased uridine supply from adipocytes, we also wondered if obesity causes a decrease in hepatic uridine synthesis and how this disruption might affect systemic energy homeostasis.

To address these questions, we examined the uridine synthesis pathway in the liver under conditions of fasting and obesity. We found the transcripts of Cad, the gene encoding carbamoyl phosphate synthetase 2, aspartate transcarbamylase, and dihydroorotase, a trifunctional enzyme for *de novo* pyrimidine synthesis [19], are decreased by fasting and diet-induced obesity. In addition, Cad downregulation is observed in the livers from *dbdb* mice, a rodent model of obesity and T2D, and in hepatocytes from individuals with NAFLD according to our analysis of online public databases [20, 21]. Moreover, Cad expression shows a circadian rhythm in the liver with its expression acutely upregulated in dark transition from light to dark. In mice, we selectively ablated hepatocyte Cad using an albumin-Cre mediated LoxP strategy. The Cad KO mice of a high fat diet (HFD) gained more weight and were more insulin resistant sooner than the control mice. The metabolic dysregulation in the KO mice was accompanied by increased hepatic gluconeogenesis and reduced hepatic insulin clearance. Interestingly, despite reduced liver uridine, the blood uridine levels remained unchanged in the KO mice under both chow and HFD feeding compared to the control mice. The maintenance of circulating uridine in the KO mice was achieved through an increase in uridine supply from the adipocytes. This is critical as we found that uridine preferentially shut down pyrimidine synthesis in hepatocytes but not adipocytes at blood levels seen in fasting. Remarkably, fasting levels of blood uridine increased hepatocyte gluconeogenesis, and this was even more so when *de novo* pyrimidine synthesis was suppressed, indicating a synergistical action of uridine and its biosynthesis pathway in promoting hepatic glucose production, which provides a new paradigm for the etiology of obesity and T2D.

## Results

### *De novo* hepatic pyrimidine biosynthesis is downregulated by HFD feeding and fasting

In obesity and diabetes, uridine plays an essential role in glycemic control to limit the development of insulin resistance (IR) [12, 22, 23]. Analysis of data in the GEO revealed a downregulation of Cad expression in the livers from *dbdb* mice, a rodent model of obesity and type 2 diabetes [20] (**Supplemental Fig. 1A**). Single-nucleus sequencing of human samples also showed a decrease in CAD expression in hepatocytes from individuals with NAFLD [21] (**Supplemental Fig. 1B**), suggesting Cad reduction is linked to obesity and T2D. To test this, we performed untargeted metabolomics study of livers harvested from C57BL/6 mice on normal chow diet (NCD) and HFD. The HFD mice showed a decrease of intermediate metabolites from *de novo* pyrimidine synthesis (orotic acid, CMP, deoxycytidine) as well as pyrimidine catabolism (dihydrothymine, β-alanine) (**Fig. 1A and Supplemental Fig. 1C**). Gene expression analysis confirmed the downregulation of Cad and Upp2 by HFD in the livers of mice (**Fig. 1B**). Liver Cad protein was reduced 40% by HFD in fed mice (**Fig. 1C**). In contrast, Cad protein in epididymal white adipose tissue (eWAT) was increased by 23% in HFD mice (**Fig. 1D**), whereas Cad protein in subcutaneous white adipose tissue (sWAT) remained unchanged (**Supplemental Fig. 1D**). To examine if the reduction in liver Cad in obesity might be triggered directly by disrupted metabolism at substrate level, we isolated hepatocytes from chow-fed WT mice and cultured the cells in serum-free media with high glucose and palmitate to mimic the metabolite changes seen in diet-induced obesity. High glucose alone or plus palmitate led to a decrease in Cad protein (**Fig. 1E**) without jeopardizing cell viability (**Supplemental Fig. 1E**), suggesting that the reduction of hepatic uridine biosynthesis could be a response to increased blood glucose and/or fatty acids in obesity. In contrast to the metabolic dysregulation in obesity, fasting, a process requiring metabolic adaptation, was reported to downregulate Cad gene expression in the liver [14]. We found that fasting for 24h caused a decrease in liver Cad mRNA (**Fig. 1F**) and protein (**Fig. 1G**), confirming that hepatic *de novo* uridine biosynthesis is downregulated by fasting. Notably, pyrimidine synthesis pathway shows a diurnal rhythms in the livers from mice on chow (**Supplemental Fig. 1F**) [24], supporting that *de novo* pyrimidine synthesis is reduced when light is on, time for mice to reduce eating. Among the pyrimidine biosynthesis genes, Cad shows the highest sensitivity to the clock (**Supplemental Fig. 1F, black line**), suggesting Cad is the key gene for this pathway to be regulated by circadian.

**Figure 1.**
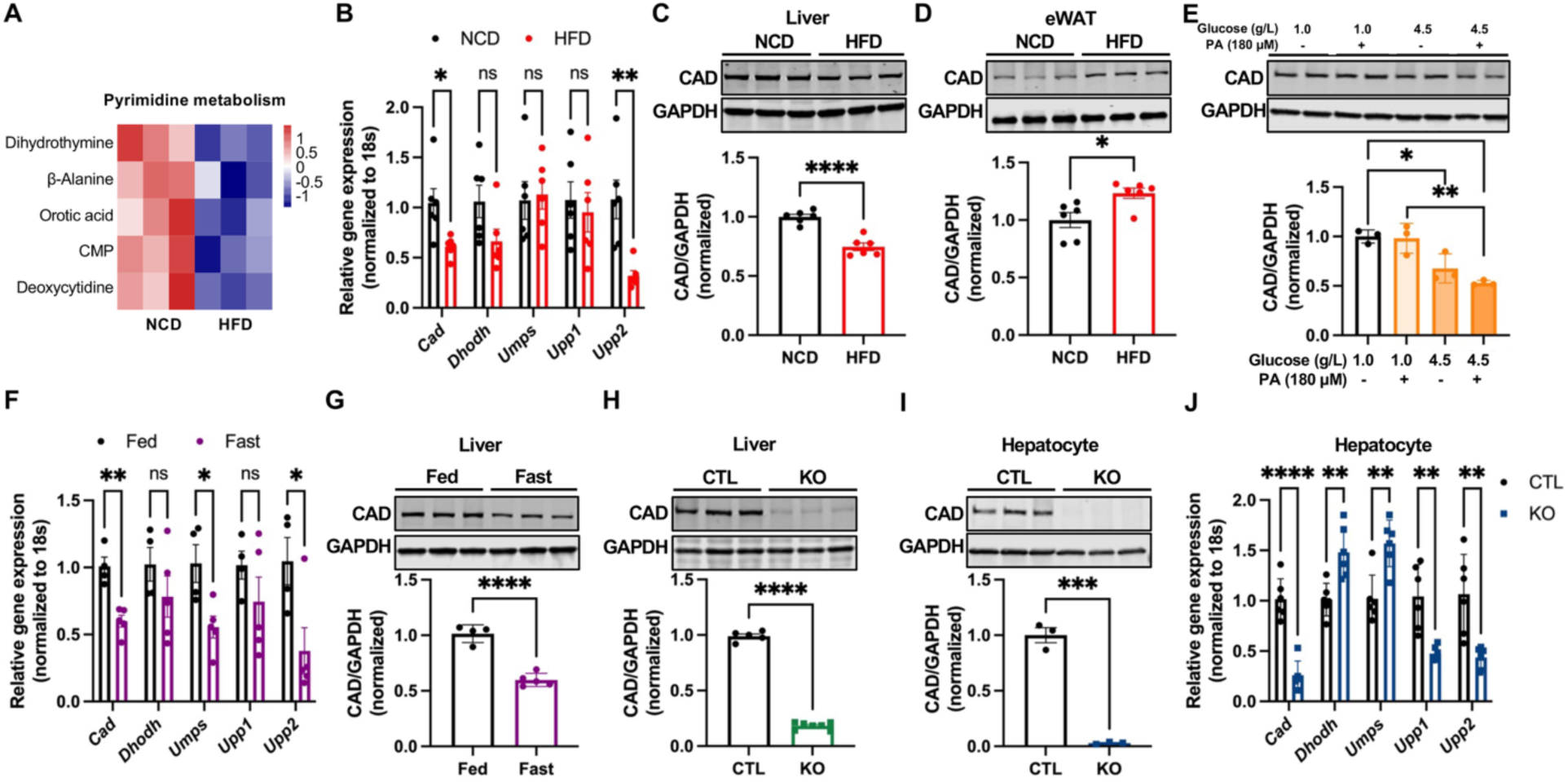
*De novo* hepatic pyrimidine biosynthesis is downregulated by HFD feeding and fasting. (**A**) Hierarchical clustering of pyrimidine metabolites in the livers from WT mice on NCD or after 15 weeks of HFD feeding (n = 3 per group). Tissues were harvested from random fed mice. (**B**) qPCR analysis of uridine metabolism genes in the livers from WT mice on NCD or after 12 weeks of HFD feeding (n = 6 per group). Tissues were harvested from random fed mice. (**C-D**) Western blot and quantification of Cad protein in the liver (**C**) and epididymal white adipose tissue (eWAT) (**D**) from WT mice on NCD or after 12 weeks of HFD feeding (n = 6 per group). Tissues were harvested from random fed mice. (**E**) Western blot and quantification of Cad protein in primary hepatocytes pretreated with glucose and palmitate as indicated for 24 hr before a 6 hr incubation in glucose production media containing 20 mM lactate and 2 mM pyruvate (n = 3 per group). (**F**) qPCR analysis of uridine metabolism genes in the livers from WT mice on NCD, fed (n = 4) or after a 24 hr fast (n = 5). (**G**) Western blot and quantification of Cad protein in the livers from WT mice on NCD, fed (n = 4) or after a 24 hr fast (n = 5). (**H**) Western blot and quantification of Cad protein in the livers from CTL (Cad^f/f^, n = 5) and KO (Cad^f/f^, Alb-Cre, n = 7) mice after 35 weeks of NCD feeding. Tissues were collected after a 24 hr fast. (**I**) Western blot and quantification of Cad protein in hepatocytes isolated from CTL and KO mice after 12 weeks of NCD feeding (n = 3 per group). (**J**) qPCR quantification of uridine metabolism genes in hepatocytes isolated from CTL and KO mice. ^*^p < 0.05, ^**^p < 0.01, ^***^p < 0.001, ^****^p < 0.0001; ns, no significance. Error bars denote SEM.

To understand the significance of hepatic uridine biosynthesis suppression in the etiology of obesity and T2D, we generated a mouse model with hepatocyte-specific deletion of Cad by crossing the Albumin-Cre mice to the Cad^flox/flox^ mice [18]. Western blot analysis demonstrated >80% reduction of Cad protein in the livers of Cad KO mice (**Fig. 1H**). Since hepatocytes make up 70% of the liver [25], the residual expression of Cad protein detected in the liver likely originates from other cell types. Isolated hepatocytes showed a complete absence of Cad protein (**Fig. 1I**), indicating an efficient ablation of hepatic Cad in the KO mice. The other two genes for *de novo* pyrimidine synthesis, Dhodh and Umps, were, however, found upregulated, and the genes for uridine catabolism, Upp1 and Upp2, were downregulated (**Fig. 1J**), suggesting that the loss of Cad causes compensatory responses in both synthesis and degradation pathways of uridine.

### Hepatic Cad loss promotes diet-induced obesity with increased lipogenesis

The decrease of hepatic Cad expression in obesity suggests that Cad loss might promote the progression of obesity. Indeed, when switched to HFD, the Cad KO mice displayed more weight gain (**Fig. 2A**) and more fat mass (**Fig. 2B**). Fasting serum cholesterol was increased in the Cad KO mice (**Fig. 2C**), while triglyceride, fatty acid, and glycerol levels remained comparable to controls (**Fig. 2C-D**). The liver content of triglyceride (**Fig. 2E**) and cholesterol, particularly esterified cholesterol (**Fig. 2F**), was much increased in the Cad KO mice. Consistent with an increased lipid accumulation, the liver-to-body weight ratio was higher for the Cad KO mice (**Fig. 2G**). Serum levels of reduced glutathione (GSH), alanine aminotransferase (ALT) and aspartate aminotransferase (AST) were significantly elevated in Cad KO mice after HFD (**Fig. 2H-I**), suggesting hepatic injury. Tissue histology revealed an increase in lipid content but not fibrosis in the livers from Cad KO mice (**Fig. 2J-K**). Notably, the Cad KO mice on NCD did not show more weight gain than the control mice when fed on NCD (**Supplemental Fig. 2A**). Also, the liver weights, blood lipid levels (except for triglyceride), and blood glycerol levels in the KO mice were all comparable to the control mice (**Supplemental Fig. 2B-F**). Serum ALT and AST levels were not elevated in the KO mice (**Fig. S2G-H**), indicating there was no liver damage in chow fed CAD KO mice. These data indicate that suppression of uridine biosynthesis in hepatocyte by Cad KO promotes diet-induced obesity although the KO *per se* does not lead to liver dysfunction in chow-fed mice.

**Figure 2.**
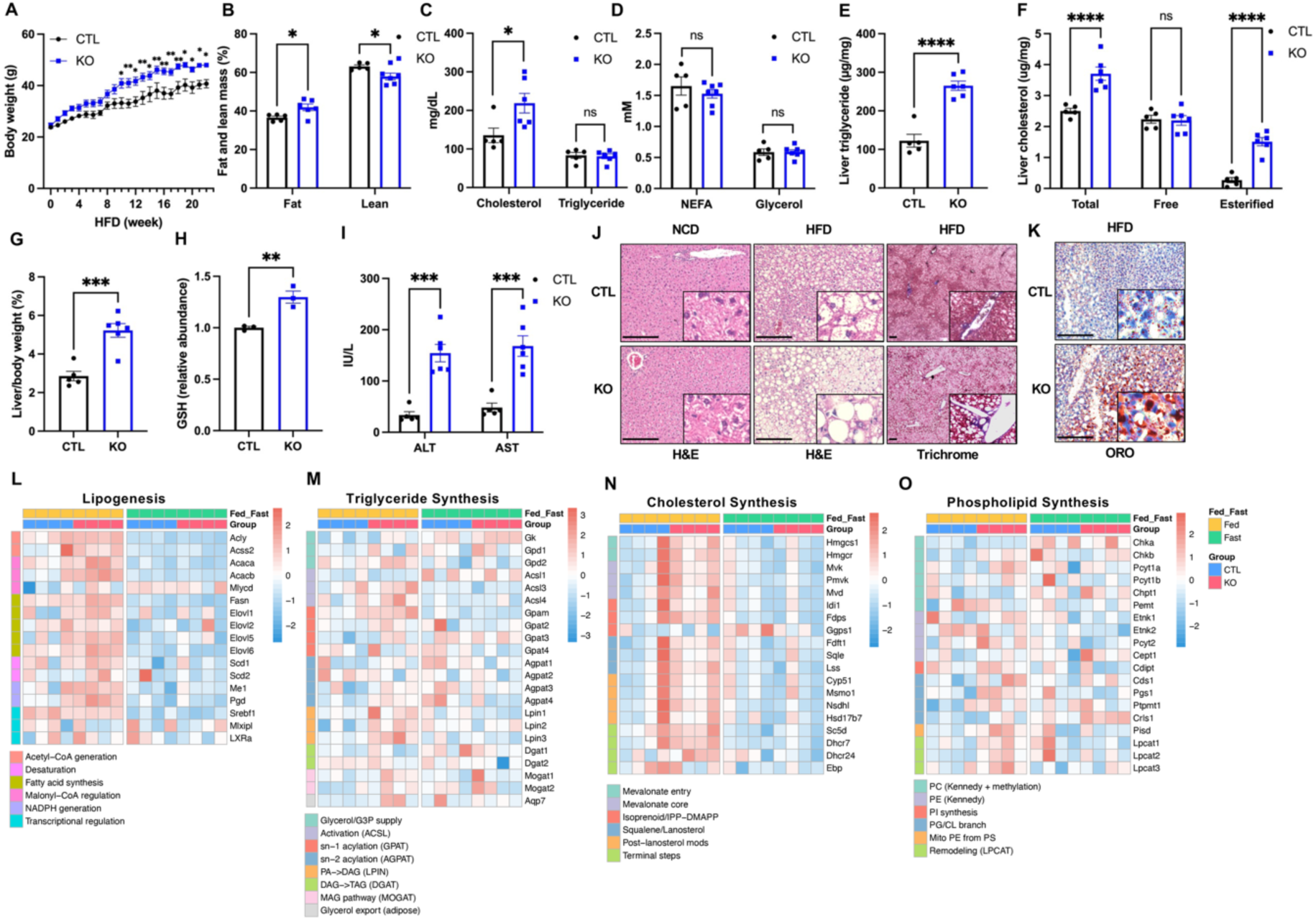
Hepatic Cad loss promotes diet-induced obesity with increased lipogenesis. (**A**) Weekly body weight for CTL (n = 5) and KO (n = 6) mice during 22 weeks of HFD feeding. (**B**) Fat and lean mass of CTL (n = 5) and KO (n = 6) mice determined by magnetic resonance imaging after 22 weeks of HFD feeding. (**C-D**) Serum cholesterol and triglyceride (**C**), NEFA and glycerol (**D**) in CTL (n = 5) and KO (n = 6) mice after 23 weeks of HFD feeding. Serum were collected after a 24 hr fast. (**E-F**) Liver triglyceride (**E**) and cholesterol (**F**) contents of CTL (n = 5) and KO (n =6) mice after 23 weeks of HFD feeding. Tissues were collected after a 24 hr fast. (**G**) Liver weight to body weight ratio of CTL (n = 5) and KO (n = 6) mice after 23 weeks HFD feeding. Measurements were done after a 24h fast. (**H**) Plasma GSH of CTL and KO mice after 23 weeks of HFD feeding. Plasma was collected after a 24 hr fast and GSH level determined by LC-MS/MS (n = 3 per group). (**I**) Plasma ALT and AST levels of CTL (n = 5) and KO (n = 6) mice after 23 weeks of HFD feeding. Plasma was collected after a 24 hr fast. (**J**) Representative hematoxylin-eosin (H&E) (under 20x objective) and trichrome staining (under 4x objective) images of the liver sections from CTL and KO mice fed on NCD or after 20 weeks of HFD feeding. Scale bar, 200 µm. Insets are magnified 16 times. Tissues were collected from random fed mice. (**K**) Representative oil red O (ORO) images of the liver cryosections from CTL and KO mice after 15 weeks of HFD feeding. Scale bar, 200 µm. Insets are magnified 16 times. Tissues were collected after a 24 hr fast. (**L-O**) Heatmap of genes in *de novo* lipogenesis (**K**), triglyceride synthesis (**L**), cholesterol synthesis (**M**), and phospholipid synthesis (**N**) pathways. Maps were generated with z-score from bulk RNA sequencing data. Liver tissues were collected from CTL and KO mice after 15 weeks of HFD feeding, fed or after a 24 hr fast (n=4 per group). ^*^p < 0.05, ^**^p < 0.01, ^***^p< 0.001, ^****^p < 0.0001; ns, no significance. Error bars denote SEM.

The increase in liver lipid in the KO mice suggests an upregulation of lipogenesis. We therefore conducted bulk RNA sequencing of the livers from HFD fed mice. The Cad KO mice showed more robust expression of genes associated with lipogenesis pathway than control mice in fed state (**Fig. 2L**). Specifically, significant upregulation of LXR⍺, Srebp1, Fasn, and Scd1 was seen in livers from KO mice even after a 24 hr fast (**Supplemental Fig. 2I**). Moreover, the synthesis pathways for TG, cholesterol, and phospholipid were all upregulated in CAD KO mice in fed state (**Fig. 2M-O**), confirming that Cad KO leads to increased lipid synthesis in hepatocytes. Notably, fasting-induced downregulation of lipogenesis and cholesterol synthesis pathways were preserved in the Cad KO mice (**Fig. 2L & 2N**), indicating that the regulation of lipid synthesis by fasting was maintained with hepatocyte Cad KO.

### Hepatic Cad loss promotes insulin resistance with concomitant reduction of insulin clearance

Obesity serves as a key predictor of insulin resistance and represents a major risk factor for T2D [26]. Since the hepatic Cad KO mice are more prone to diet-induced obesity, they are expected to develop insulin resistance more severely than the control group. To trace the progression of insulin resistance on HFD, we conducted oral glucose tolerance tests (OGTT) every 5 weeks. At the end of the first 5-week-HFD feeding interval, no difference was detected in blood glucose levels between Cad KO and control mice in OGTT (**Fig. 3A-B**). After 10 and 15 weeks of HFD feeding, CAD KO mice exhibited a trend toward impaired glucose tolerance (**Fig. 3A-B**). In contrast, a significant elevation in glucose-stimulated insulin was observed in the Cad KO mice during each OGTT (**Fig. 3C-D**), indicating impaired insulin sensitivity in the KO mice. By the end of 20 week of HFD, fasting glucose (**Fig. 3E**), fasting insulin (**Fig. 3F**), and HOMA-IR (**Fig. 3G**) were all higher in the CAD KO mice, suggesting the presence of a more insulin resistance. Hyperinsulinemic-euglycemic clamp study were carried out to more fully assess this (**Fig. 3H**). The glucose infusion rate (**Fig. 3I**) and total glucose infused (**Fig. 3J**) were lower in the Cad KO mice which was associated with reduced glucose uptake in the tibialis anterior muscle (**Fig. 3K**). The clamp data indicates a more severe insulin resistance in the Cad KO mice by 18 weeks of HFD feeding. Interestingly, when fed on NCD, the Cad KO mice showed no difference in glycemic control and insulin sensitivity compared to control group including a fed/fast/refed study (**Supplemental Fig. 3A**), OGTT (**Supplemental Fig. 3B-E**), and hyperinsulinemic-euglycemic clamp (**Supplemental Fig. 3F-I**).

**Figure 3.**
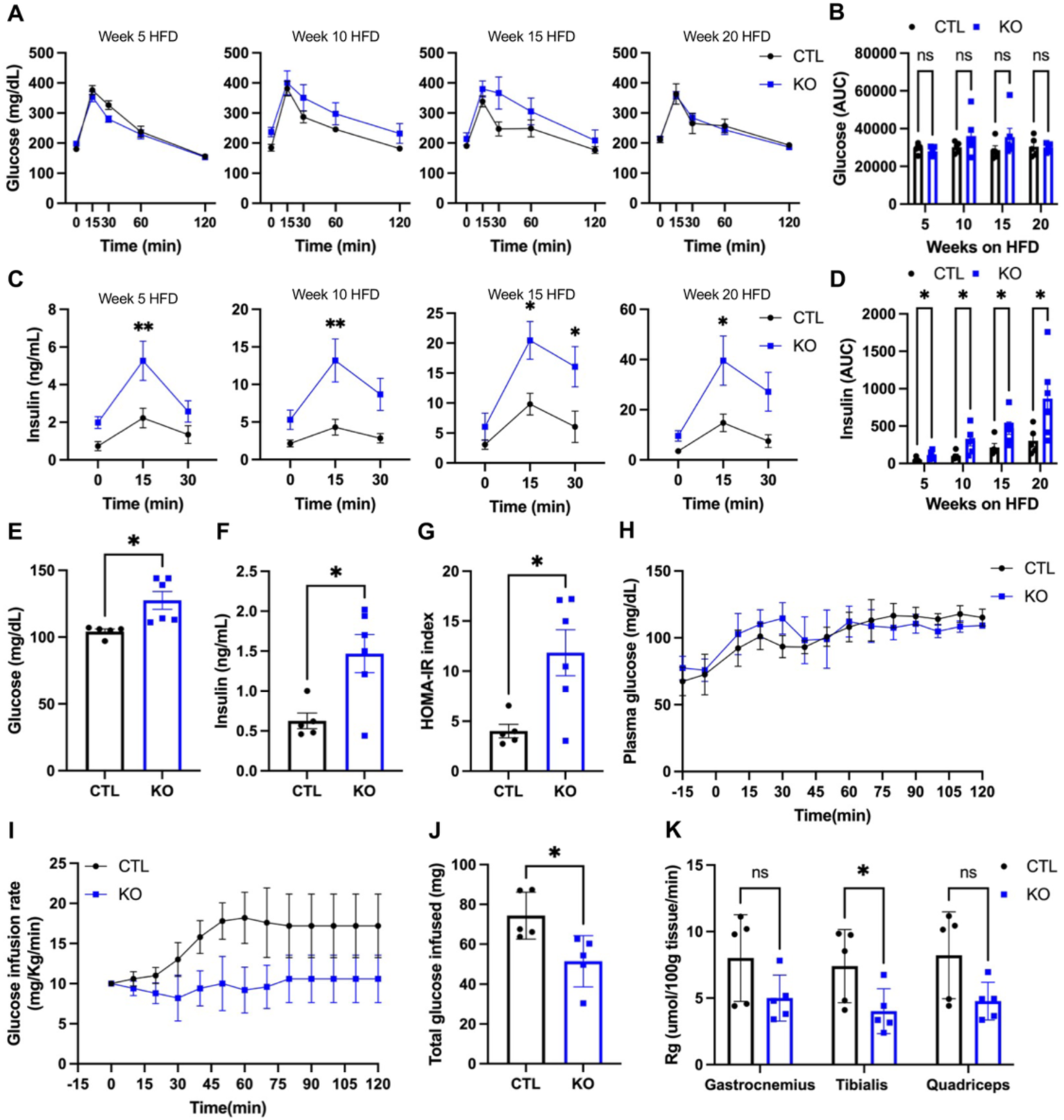
Hepatic Cad loss exacerbates insulin resistance in diet-induced obesity. (**A-B**) Blood glucose response to glucose overload (**A**) and the area under curve (AUC) (**B**) in oral glucose tolerance tests (OGTTs) in CTL (n = 5) and KO (n = 6) mice during HFD feeding. (**C-D**) Plasma insulin response to glucose overload (**C**) and the AUC (**D**) in the OGTTs in (**A**). (**E-F**) Fasting blood glucose (**E**) and insulin levels (**F**) in CTL (n = 5) and KO (n = 6) mice after 23 weeks of HFD feeding. Plasma was collected after a 24 hr fast. (**G**) HOMR-IR was calculated from the data in **E** and **F** using the formula FBG✕FINS/22.5. (**H**) Blood glucose levels at 15-minute intervals throughout hyperinsulinemic-euglycemic clamp for CTL and KO mice after 18 weeks of HFD feeding (n = 5 each group). (**I-J**) Glucose infusion rate (**I**) and total glucose infused (**J**) during the hyperinsulinemic-euglycemic clamp study in (**H**). (**K**) Gastrocnemius, tibialis anterior, and quadriceps muscle glucose uptake during the hyperinsulinemic-euglycemic clamp study in (**H**). ^*^p < 0.05, ^**^p < 0.01; ns, no significance. Error bars denote SEM.

In addition to insulin resistance which drives insulin hypersecretion, reduced insulin clearance also contributes to the development of hyperinsulinemia [27, 28]. About 50% insulin in the portal vein is degraded in hepatocytes, a process known as first-passage insulin clearance [29]. To examine the impact of hepatic Cad loss on hepatic insulin clearance, we measured the response of plasma insulin and C-peptide to glucose overload in the Cad KO mice after 10wk of HFD feeding. The acute elevation of insulin and C-peptide by glucose overload was higher in Cad KO mice than the control group (**Fig. 4A-D**). However, the ratio of C-peptide-to-insulin was lower in Cad KO mice (**Fig. 4E-F**), suggesting a decrease in hepatic insulin clearance in the Cad KO mice. Carcinoembryonic antigen-related cell adhesion molecule 1 (CEACAM1), a substrate of the insulin receptor in the liver, promotes hepatic insulin clearance [30]. We found CEACAM1 was significantly reduced in livers from Cad KO mice (**Fig. 4G-H**). Thus, hepatocyte Cad loss appears to suppress insulin clearance in the liver by downregulation of CEACAM1. A decrease in hepatic insulin clearance could potentially protect islet β-cells from exhaustion due to insulin hypersecretion triggered by insulin resistance. Consistent with this idea, the islet architecture of pancreata from Cad KO mice was not abnormal (**Supplemental Fig. 4A**). Immunofluorescent staining of pancreatic sections also suggest that insulin-producing β-cells and glucagon-producing ɑ-cells were comparable between the KO and control samples (**Supplemental Fig. 4B**).

**Figure 4.**
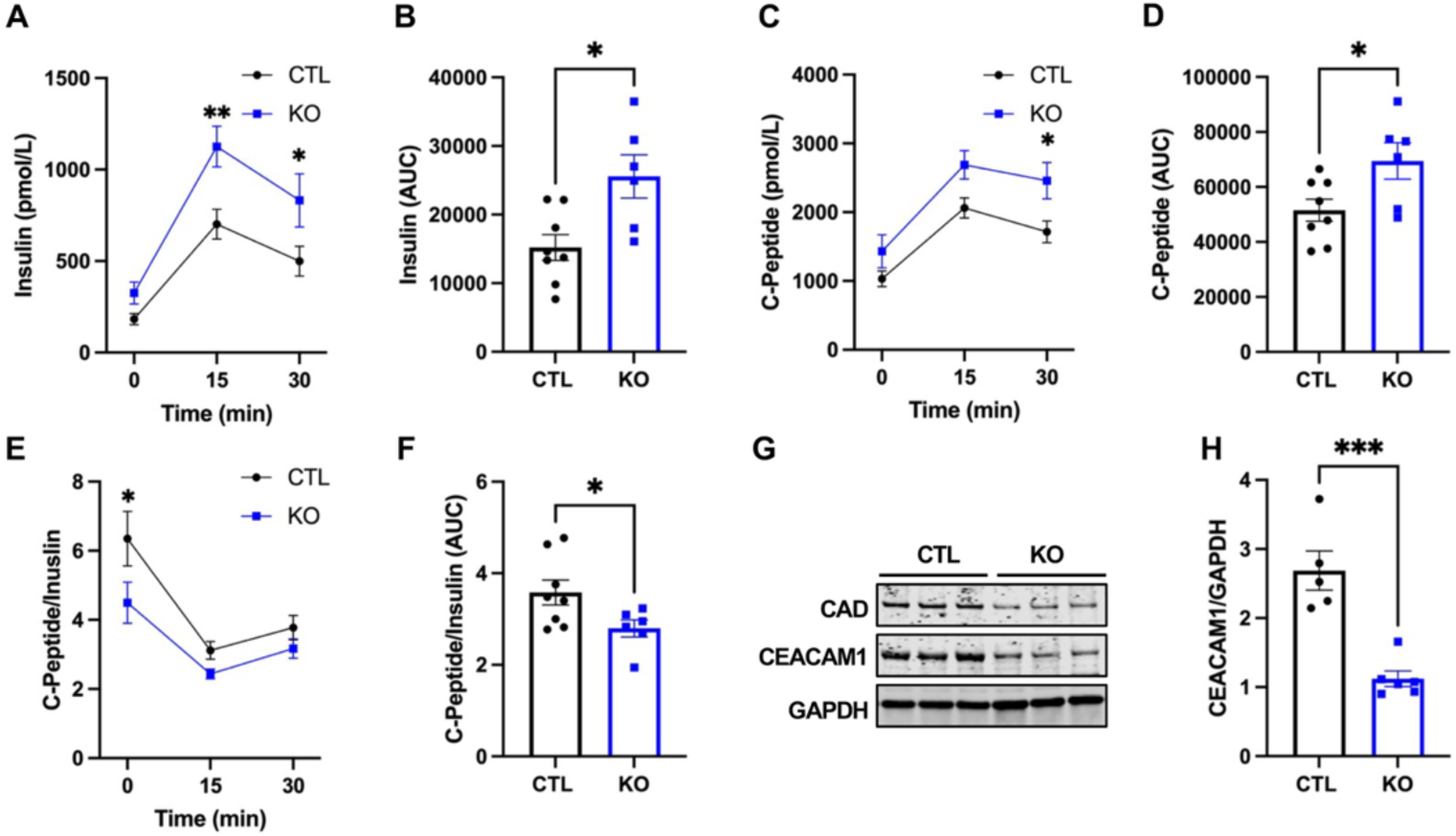
Hepatic Cad KO reduces insulin clearance in diet-induced obesity. (**A-B**) Plasma insulin response to glucose overload (**A**) and the AUC (**B**) during OGTT in CTL (n = 8) and KO (n = 6) mice after 10 weeks of HFD feeding. (**C-D**) Plasma C-peptide response to glucose overload (**C**) and the AUC (**D**) during the OGTT in (**A**). (**E-F**) C-peptide-to-insulin molar ratio response to glucose overload (**E**) and the AUC (**F**) during the OGTT in (**A**). (**G-H**) Western blot (**G**) and quantification (**H**) of Ceacam1 protein in the livers from CTL (n = 5) and KO (n = 6) mice after 23 weeks of HFD feeding. Tissues were collected after a 24h fast. ^*^p < 0.05, ^**^p < 0.01, ^***^p< 0.001. Error bars denote SEM.

### Enhanced lipid oxidation in hepatic Cad KO mice fuels gluconeogenesis

Despite of the presence of more severe insulin resistance and obesity, the Cad KO mice on HFD did not show an elevation in fasting FFA (**Fig. 2D**), suggesting fatty acid oxidation is enhanced and/or TG mobilization is decreased. To assess this, metabolic cage studies were done. The Cad KO mice exhibited a significantly lower respiratory exchange ratio (RER) during dark and light intervals (**Fig. 5A and 5C**), supporting a shift in fuel usage from carbohydrates to lipids [31]. The Cad KO mice also showed a reduction in energy expenditure (**Fig. 5B and 5D**) despite that their energy intake (**Fig. 5E**) and locomotive activity (**Fig. 5F**) were not different from the control group, indicating their higher rates of weight gain result partially from reduced energy expenditure.

**Figure 5.**
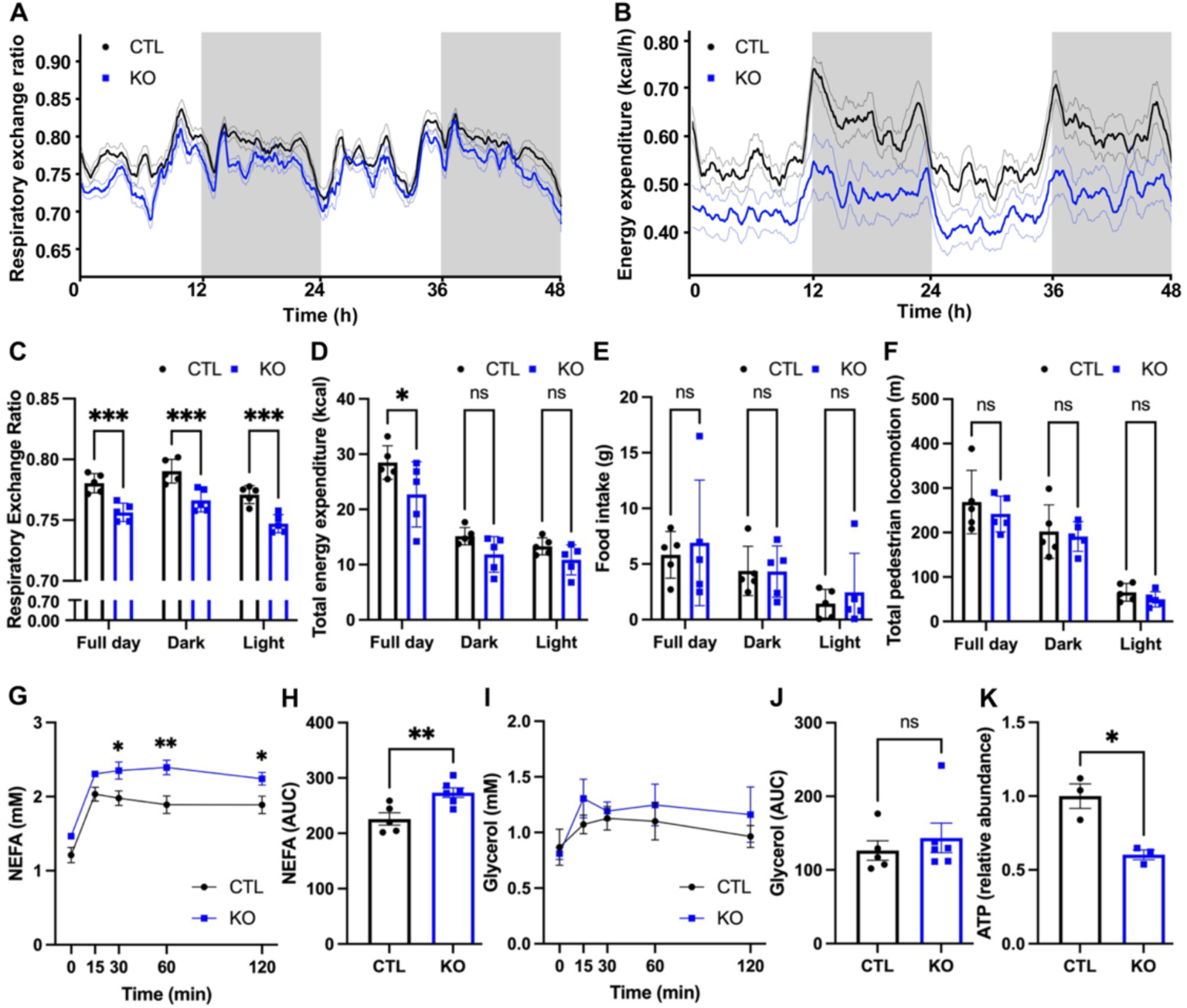
Impact of hepatic Cad KO on systemic energy homeostasis. (**A-F**) Respiratory exchange ratio (**A & C**), energy expenditure (**B & D**), food intake (**E**), and locomotion (**F**) in CTL and KO mice after 15 weeks of HFD feeding (n = 5 per group). (**G-J**) Serum NEFA and glycerol response to CL316,243 (**G & I**) and the AUCs (**H & J**) in CTL (n = 5) and KO (n = 6) mice after 18 weeks of HFD feeding. (**K**) Liver ATP levels in mice after 15 weeks of HFD feeding. Tissues were collected after a 24 hr fast and ATP determined by LC-MS/MS (n = 3 per group). ^*^p < 0.05, ^**^ p < 0.01, ^***^ p < 0.001; ns, no significance. Error bars denote SEM.

To examine if TG mobilization by fasting is reduced in the Cad KO mice, we administered to mice CL316,243, a selective β3-adrenergic receptor agonist [32, 33] that stimulates adipocyte lipolysis. The treated Cad KO mice showed a higher release of FFA than the controls (**Fig. 5G-H**). The higher rate of FFA release observed in Cad KO mice is consistent with the observation that the Cad KO mice have more fat (**Fig. 2B**). Interestingly, in the KO mice, CL316,243 only moderately increased glycerol release (**Fig. 5I-J**), suggesting that the consumption of glycerol is higher in the KO mice. Notably, the ATP levels were significantly reduced in the liver of Cad KO mice (**Fig. 5K**). Therefore, the metabolic remodeling in the Cad KO mice is accompanied by increased FFA oxidation relative to glucose, increased glycerol consumption, and decreased hepatic ATP level. This remodeling could potentially result from increased gluconeogenesis in hepatocytes [34]. Consistent with this idea, we found that precursors for gluconeogenesis were elevated in the blood from fasted Cad KO mice (**Fig. 6A** & **Supplemental Fig. 5A**). Indeed, the gluconeogenic pathway was significantly altered in the livers of Cad KO mice (**Fig. 6B**). As expected, the beta oxidation and gluconeogenesis pathways were upregulated in the Cad KO mice in fed state when examined by RNA bulk sequencing (**Supplemental Fig. 5B-C**).

**Figure 6.**
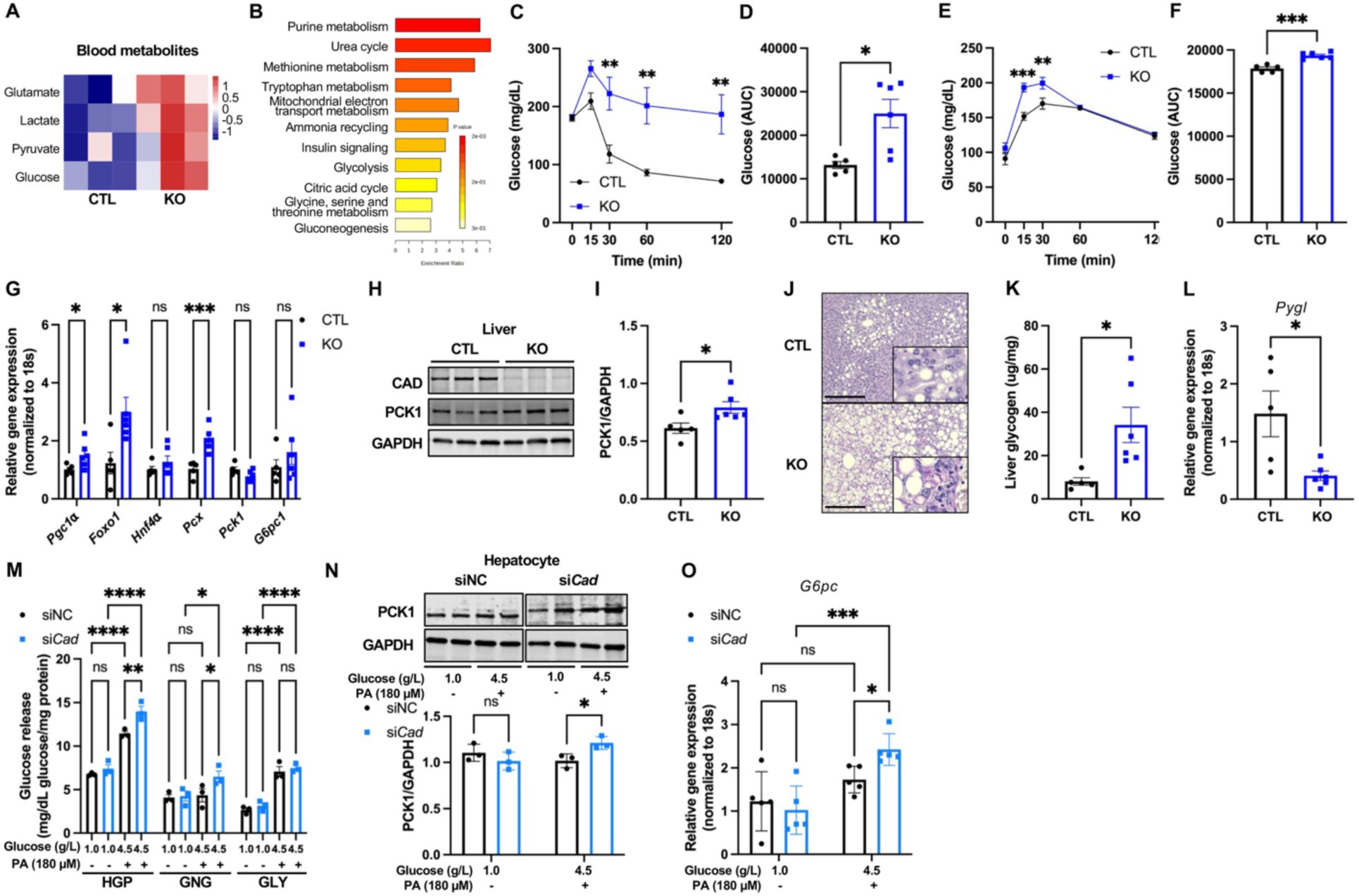
Enhanced lipid oxidation in hepatic Cad KO mice fuels gluconeogenesis. (**A**) Hierarchical clustering of blood metabolites from CTL and KO mice after 23 weeks of HFD feeding. Plasma were collected after a 24 hr fast and metabolites determined by LC-MS/MS (n = 3 per group). (**B**) Metabolite sets enrichment in the livers by Cad KO. Tissues were collected from CTL and KO mice after 15 weeks of HFD feeding. Metabolites were determined by LC-MS/MS and analyzed with MetaboAnalyst 6.0. (n = 3 per group). (**C-D**) Blood glucose response to CL316,243 (**C**) and the AUC (**D**) in CTL (n = 5) and KO (n = 6) mice after 18 weeks of HFD feeding. (**E-F**) Blood glucose response to pyruvate (**E**) and the AUC (**F**) in CTL (n = 5) and KO (n = 6) mice after 6 weeks of HFD feeding. (**G**) qPCR analysis of gluconeogenic genes in the livers from CTL (n = 5) and KO (n = 6) mice after 23 weeks of HFD feeding. Tissues were collected after a 24h fast. (**H-I**) Western blot (**H**) and quantification (**I**) of PCK1 protein in the livers from CTL (n = 5) and KO (n = 6) mice after 23 weeks of HFD feeding. Tissues were collected after a 24 hr fast. (**J**) Representative PAS staining of liver sections from CTL and KO mice after 15 weeks of HFD feeding. Scale bar, 200 µm. Insets are magnified 16 times. Tissues were collected after a 24 hr fast. (**K**) Liver glycogen contents in CTL (n = 5) and KO (n = 6) mice after 23 weeks of HFD. Tissues were collected after a 24 hr fast. (**L**) qPCR analysis of Pygl in the livers from CTL (n = 5) and KO (n = 6) mice after 23 weeks of HFD. Tissues were collected after a 24 hr fast. (**M**) Hepatic glucose production (HGP), gluconeogenesis (GNG), and glycogenolysis (GLY) in WT primary hepatocytes after siNC or si*Cad*-mediated knockdown. The hepatocytes received a 24 hr exposure to glucose and palmitate as indicated before the 6 hr incubation in glucose production media that contained 20 mM lactate and 2 mM pyruvate. (n = 3 per group). (**N**) Western blot and quantification of PCK1 protein in hepatocytes after the siNC or si*Cad*-mediated knockdown in (**M**) (n = 3 per group). (**O**) qPCR analysis of G6pc in WT primary hepatocytes after the siNC or si*Cad*-mediated knockdown and 24 hr exposure to glucose and palmitate as indicated (n = 3 per group). ^*^*p* < 0.05, ^**^*p*< 0.01, ^***^*p*< 0.001, ^****^*p*< 0.0001; ns, no significance. Error bars denote SEM.

To test if gluconeogenesis is increased in Cad KO mice, the impact of CL316,243 on blood glucose is assessed. In fasted mice, the FFA released from adipocytes via lipolysis is used by hepatocytes as a fuel to power gluconeogenesis. CL316,243 causes an acute increase in circulating insulin [35]. Therefore, while CL316,243 induces lipolysis, the increased gluconeogenesis does not always cause a detectable elevation in blood glucose. Thus, after CL316,243, mice may exhibit a fall in blood glucose instead of an elevation. Since HFD feeding impairs insulin sensitivity in mice, a transient elevation in blood glucose by CL316,243 might occur. This is exactly what we observed in the control HFD group (**Fig. 6C**, black line at 15 minutes). CL316,243 stimulated a bigger elevation in blood glucose in KO than the controls (**Fig. 6C**, blue line at 15 minutes). Moreover, the elevation in blood glucose in the CAD KO mice was maintained for 2 hours whereas in the controls blood glucose dropped by 60% (**Fig. 6C**). Area under curve analysis confirmed that the circulating glucose was significantly higher in the Cad KO mice than the controls after CL316,243 (**Fig. 6D**). As expected, CL316,243 increased circulating insulin levels in both groups (**Supplemental Fig. 5D-E**). Similar to glucose-induced insulin secretion in OGTT (**Fig. 3C-D**), the Cad KO mice showed a higher release of insulin in response to CL316,243 than the control mice. Thus, stimulated-lipolysis in Cad KO mice is associated with higher glucose production from the liver.

The pyruvate tolerance test (PTT), by supplying exogenous gluconeogenetic substrate, directly assesses the difference in gluconeogenesis *in vivo*. Pyruvate elevated blood glucose more in Cad KO mice than controls (**Fig. 6E-F**), again indicating that hepatic Cad KO leads to increased gluconeogenesis. Pgc1⍺, a master regulator of FFA beta oxidation, and Foxo1 and Pcx, two key gluconeogenic genes were found upregulated at the transcript level in the livers from the KO mice (**Fig. 6G**), whereas phosphoenolpyruvate carboxykinase 1 (PCK1), the rate-limiting gluconeogenic enzyme that is regulated by insulin [36], was elevated at the protein level (**Fig. 6H-I**).

Increased gluconeogenesis may not lead to frank diabetes as the glucose can be used by anabolic pathways. Glucose-6-phosphatase (G6pc) converts glucose-6-phosphate to glucose for release into the blood. In livers from Cad KO mice, expression of G6pc was not increased proportionally to Foxo1 or Pcx (**Fig. 6G**), suggesting that the hepatic release of glucose to the blood might be restricted and that the glucose produced from augmented gluconeogenesis could be shunned to anabolic pathways. Indeed, hepatic glycogen content was found significantly elevated in Cad KO mice (**Fig. 6J-K**). Moreover, Glycogen phosphorylase (Pygl) was found downregulated at transcript level (**Fig. 6L**), suggesting that in Cad KO mice carbohydrate metabolism is rewired to favor both gluconeogenesis and glycogen storage. The metabolic remodeling by CAD KO was specific to HFD as no change in hepatic gluconeogenic capacity (**Supplemental Fig. 5F-G**) or hepatic glycogen content (**Supplemental Fig. 5H-I**) was detected in the Cad KO mice fed on NCD.

To examine whether the increase in gluconeogenesis by HFD in Cad KO mice is cell autonomous rather than do to insulin resistance or hepatic lipid buildup, Cad was knocked down with small interference RNAs (si*Cad*) in hepatocytes isolated from chow-fed WT mice (**Supplemental Fig. 6A-B**). The metabolic stress of obesity was mimicked by 24h exposure to high glucose plus high palmitate (HGHP, 24 hr) which did not cause cell death (**Supplemental Fig. 6C**). Hepatic glucose production was significantly elevated by HGHP regardless of Cad knockdown (**Fig. 6M, HGP**), suggesting a HGHP-dependent increase in hepatic glucose production. The Cad knockdown did not change basal hepatic glucose production (**Fig. 6M**, **HGP**, 1.0 g/L glucose & 0 μM PA). However, it caused a further elevation in glucose production in the hepatocytes pretreated with HGHP (**Fig. 6M, HGP**). The total hepatic glucose production was attributed to gluconeogenesis and glycogenolysis. Gluconeogenesis was increased by HGHP only in the hepatocytes with Cad knockdown (**Fig. 6M, GNG**), while glycogenolysis was increased by HGHP equally in the Cad knockdown and control hepatocytes (**Fig. 6M, GLY**). HGHP increased PCK1 protein in the Cad knockdown hepatocytes (**Fig. 6N**), similar to what is observed in the livers from the Cad KO mice (**Fig. 6H-I**), supporting that gluconeogenesis is increased by Cad knockdown. Also, HGHP increased G6pc mRNA in the Cad knockdown hepatocytes by (**Fig. 6O**) which was different from what was observed in the liver (**Fig. 6G**), suggesting that the glucose release in cultured hepatocytes with Cad knockdown is more robust than that in the liver of Cad KO mice. The *in vitro* results agree with the *in vivo* ones in supporting that loss or reduction of Cad in hepatocytes promotes gluconeogenesis in diet-induced obesity.

### Impact of hepatic Cad loss on hepatic uridine production and circulating uridine

Hepatocyte-specific KO of CAD is expected to abolish *de novo* pyrimidine synthesis in the liver, which will reduce a major uridine supply to blood. Since uridine has multifaceted functions, it was not clear if the metabolic sequala in Cad KO mice was mediated through disrupted uridine homeostasis in the blood. We first examined if Cad KO is sufficient to reduce uridine release from hepatocytes. Mouse primary hepatocytes were cultured in uridine-free DMEM medium in the absence of serum (**Supplemental Fig. 7A**). The Cad KO hepatocytes released less uridine into the medium after 24h culture (**Fig. 7A**). Notably, the uridine release from control hepatocytes was reduced by HGHP pretreatment, whereas the Cad KO hepatocytes were not affected by HGHP (**Fig. 7A**), suggesting that the reduction of uridine release by HGHP was mediated through suppression of *de novo* pyrimidine synthesis. To confirm that reduced Cad activity lowers uridine release, Cad function was disrupted with N-(phosphonacetyl)-Laspartate (PALA), a pharmacological Cad inhibitor [37], and Cad siRNA knockdown. Both approaches revealed that hepatic uridine release was reduced by suppression of Cad (**Fig. 7B-C**). The doses of PALA used in the study did not affect cell viability (**Supplemental Fig. 7B**).

**Figure 7.**
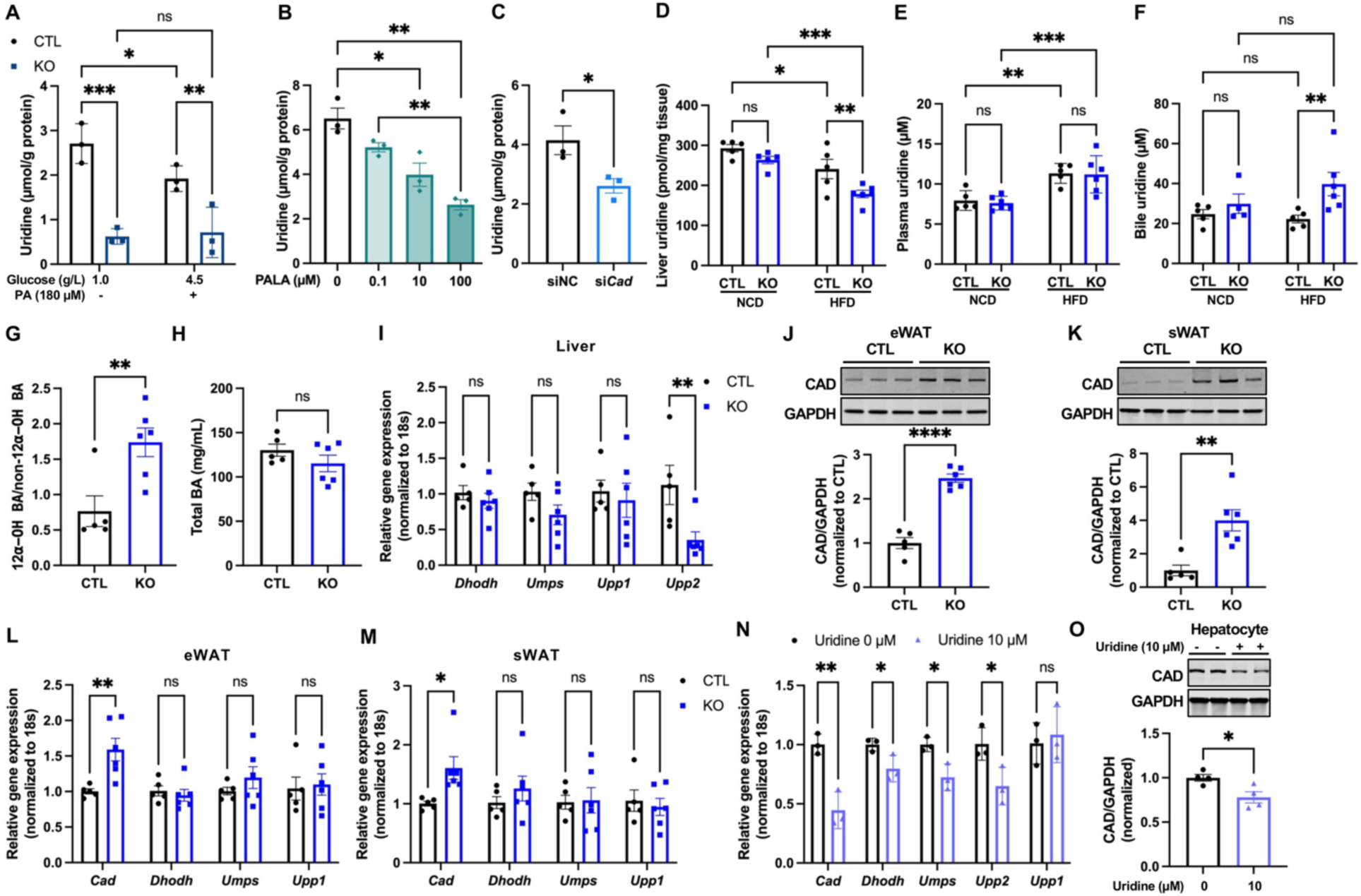
Impact of hepatic Cad KO on hepatic uridine production and circulating uridine levels. (**A**) Uridine levels in the cell culture media of primary hepatocytes isolated from CTL and KO mice following a 24 hr exposure to glucose and palmitate as indicated (n = 3 per group). (**B**) Uridine levels in the cell culture media of WT primary hepatocytes exposed to different dose of PALA for 24 hr (n = 3 per group). (**C**) Uridine levels in the cell culture media of WT primary hepatocytes after siNC or si*Cad*-mediated knockdown (n = 3 per group). (**D**) Liver uridine contents of CTL and KO mice on NCD or after 23 weeks of HFD feeding. Tissues were collected after a 24 hr fast (n = 5 ∼ 6 per group). (**E**) Plasma uridine levels of CTL and KO mice on NCD or after 16 weeks of HFD feeding. Plasma was collected after a 24 hr fast (n = 5 ∼ 6 per group). (**F**) Biliary uridine of CTL and KO mice on NCD or after 23 weeks of HFD feeding (n = 4 ∼ 6 per group). Bile was collected from gallbladder after a 24h fast. (**G-H**) Ratio of 12α-hydroxylated bile acids to non-12α-hydroxylated bile acids (**G**) and total bile acid (**H**) in the bile of CTL and WT KO mice after 23 weeks of HFD feeding (n = 4 ∼ 6 per group). Bile was collected from gallbladder after a 24 hr fast. (**I**) qPCR analysis of genes for uridine metabolism in the livers from CTL (n = 5) and KO (n = 6) mice after 23 weeks of HFD feeding. Tissues were collected after a 24 hr fast. (**J-K**) Western blot and quantification of Cad protein in the epididymal white adipose tissue (eWAT) and subcutaneous white adipose tissue (sWAT) from CTL (n = 5) and KO (n = 6) mice after 23 weeks of HFD feeding. Tissues were collected after a 24 hr fast. (**L-M**) qPCR analysis of uridine metabolism genes in the eWATs and sWATs from CTL (n = 5) and KO (n = 6) mice after 23 weeks of HFD feeding. Tissues were collected after a 24 hr fast. (**N**) qPCR analysis of uridine metabolism genes in WT primary hepatocytes after 24 hr exposure to 10 µM uridine supplemented to serum-free DMEM (n = 3 per group). (**O**) Western blot and quantification of Cad protein in primary hepatocytes after 24 hr exposure to 10 µM uridine supplemented to serum-free DMEM (n = 4 per group). ^*^p < 0.05, ^**^p < 0.01, ^***^p< 0.001, ^****^p< 0.0001; ns, no significance. Error bars denote SEM.

Liver uridine content was significantly reduced by HFD in both control and Cad KO mice, suggesting an HFD-dependent decrease in liver uridine (**Fig. 7D**). This is consistent with our finding that Cad expression is downregulated by HFD in WT mice (**Fig. 1B-C**). Liver uridine content in Cad KO mice did not show a difference from control mice when fed on NCD. However, HFD caused a 30% reduction of liver uridine content in CAD KO mice but only 18% in control mice (**Fig. 7D**), indicating that hepatocyte Cad KO aggravates the loss of liver uridine by HFD. Similar effects were observed in female hepatocyte-selective Cad KO mice (**Supplemental Fig. 7C**).

In contrast to liver uridine content, fasting blood uridine was significantly increased by HFD in both control and Cad KO mice (**Fig. 7E**). Surprisingly, the Cad KO mice maintained their fasting blood uridine at the same levels as the controls regardless of diet (**Fig. 7E**). HFD caused a 43% increase of fasting blood uridine in the control mice and 47% in the Cad KO mice (**Fig. 7E**), suggesting loss of hepatic Cad is not sufficient to prevent HFD-induced elevation in fasting uridine. Different from male mice, female mice showed no elevation in fasting uridine by HFD or by hepatocyte Cad KO (**Supplemental Fig. 7D**). Since hepatic uridine is also released to the gallbladder as a bile metabolite [14], we measured uridine level in bile. Bile uridine levels were not different between Cad KO and control mice when fed on NCD (**Fig. 7F**). However, after HFD, bile uridine levels were significantly higher in the male Cad KO mice (**Fig. 7F**). In contrast, female Cad KO mice showed increased bile uridine in NCD and after HFD (**Supplemental Fig. 7E**). Thus, hepatic Cad KO, unexpectedly, does not cause a change in circulating uridine. Instead, the liver uridine content and bile uridine levels are changed reciprocally by hepatic CAD KO. The change in bile uridine might alter bile acid composition. Indeed, the Cad KO mice showed an increase in the ratio of 12⍺-hydroxylated bile acid to non 12⍺-hydroxylated bile acids despite no change in total bile acid (**Fig. 7G-H**), suggesting an increase in bile acid-facilitated lipid absorption from foods in the Cad KO mice [38].

Dhodh and Umps, two additional genes encoding enzymes for uridine synthesis downstream of Cad, were upregulated, whereas Upp1 and Upp2, two genes encoding enzymes for uridine catabolism, were downregulated in the Cad KO hepatocytes from chow fed group (**Fig. 1J**), suggesting Cad KO triggers compensatory response in both uridine synthesis and degradation. In contrast to chow fed groups, no such compensatory changes were detected for Dhodh, Umps, and Upp1 in the livers from Cad KO mice fed the HFD (**Fig. 7I**). Upp2, the hepatocyte-specific isoform of uridine phosphorylase [39], however, showed a 68% decrease in Cad KO mice (**Fig. 7I**), which is consistent with the findings in the chow fed groups (**Fig. 1J**) except for being further reduced. Thus, HFD prevents the compensatory response to hepatic Cad loss except for Upp2.

In addition to the liver, blood uridine is also supplied by adipocytes in mice [14]. Then the diminished uridine supply from hepatocytes in the Cad KO mice might trigger a compensatory increase of uridine from their adipose tissue. Indeed, an over 2-fold increase of Cad protein was observed in eWATs and sWATs from the hepatocyte-selective Cad KO mice fed on a HFD (**Fig. 7J-K**). This was associated with 1.6-fold increase in Cad gene expression (**Fig. 7L-M**). These data suggests that circulating uridine might act as a metabolite signal to coordinate uridine production from the liver and adipose tissue. This model predicts hepatocytes would adjust their rate of uridine biosynthesis according to the circulating uridine levels. To test it, primary hepatocytes were cultured in medium with uridine (10 μM) the blood uridine level noted in fasting mice [14]. *De novo* pyrimidine synthesis genes were downregulated by 24 hr (**Fig. 7N**), while Upp2 transcripts were reduced rather than increased (**Fig. 7N**), indicating uridine suppresses at the same time uridine biosynthesis and catabolism in hepatocytes. Consistent with the decrease in Cad transcripts, the CAD protein was also reduced by uridine (**Fig. 7O**).

### Uridine synergistically increases hepatic glucose production with hepatocyte Cad loss

Since uridine causes a downregulation of *de novo* pyrimidine synthesis in hepatocytes, uridine might promote gluconeogenesis through suppressing pyrimidine synthesis. This idea was tested in cultured hepatocytes using various substrates for gluconeogenesis. When uridine was omitted, HGHP-driven gluconeogenesis was more robust in Cad KO hepatocytes than control cells in media supplemented with glutamine/pyruvate/lactate (**Fig. 8A**). Cad siRNA knockdown hepatocytes also had higher rates of gluconeogenesis than control cells in media supplemented with glutamine/pyruvate/lactate (**Fig. 8B**). Notably, glutamine-facilitated gluconeogenesis was increased by Cad loss regardless of HGHP pretreatment (**Fig. 8A-B, GNG**), which is different from the pyruvate and lactate-facilitated gluconeogenesis (**Fig. 6M**), suggesting that when Cad is suppressed the presence of glutamine can sensitizes hepatocytes to gluconeogenesis. The Cad KO hepatocytes did not show an increase in gluconeogenesis when the substrates were pyruvate and lactate (**Fig. 8C**), which is different from the hepatocytes with Cad knockdown. However, when 10 µM uridine was added the Cad KO hepatocytes pretreated with HGHP showed a significant increase in glucose production via gluconeogenesis (**Fig. 8D-F**), indicating uridine promotes gluconeogenesis synergistically with Cad loss in hepatocytes.

**Figure 8.**
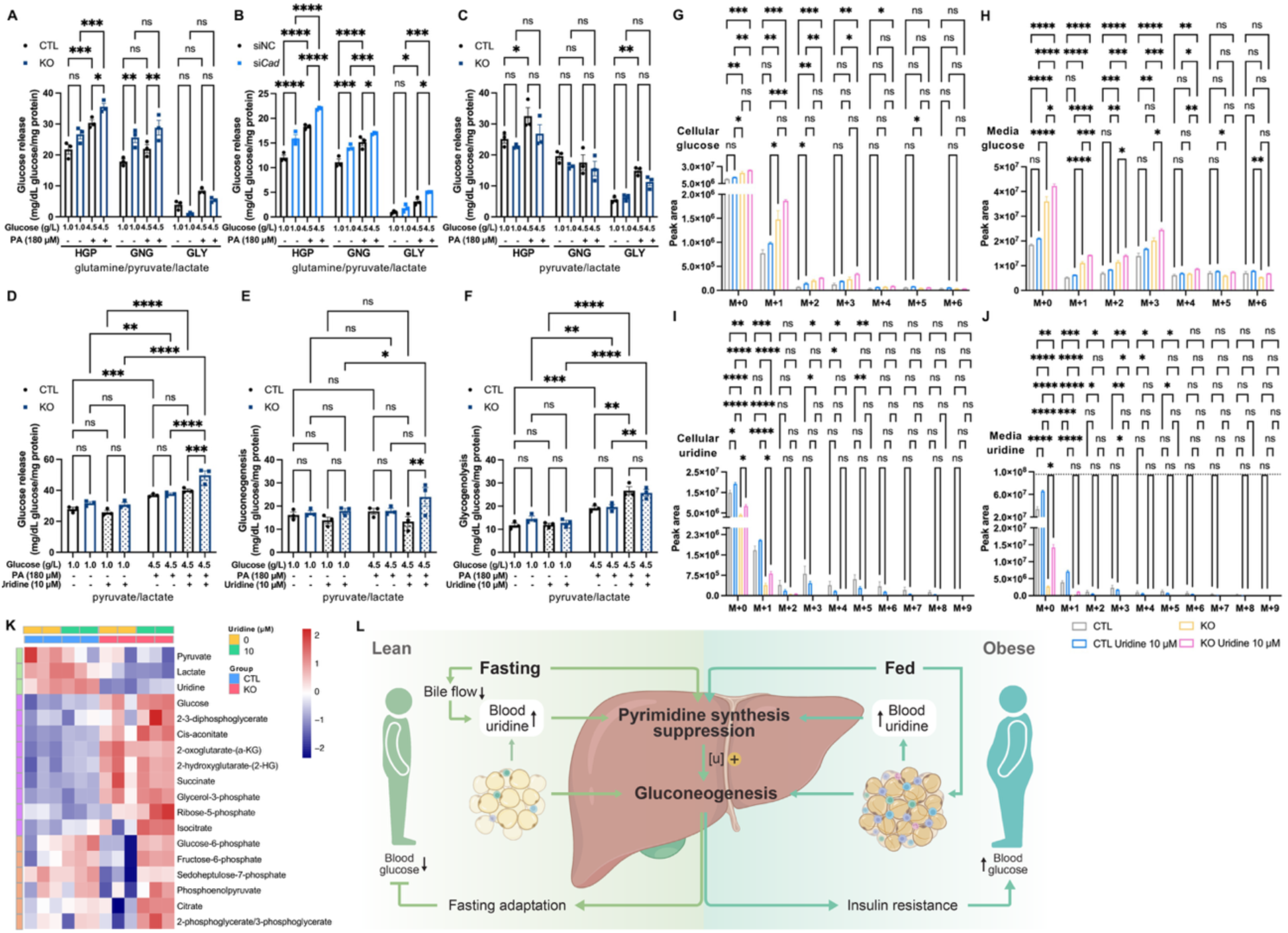
Uridine acts synergistically with hepatocyte Cad loss to increase gluconeogenesis. (**A-B**) Hepatic glucose production (HGP), gluconeogenesis (GNG), and glycogenolysis (GLY) in primary hepatocytes from CTL and KO mice (**A**), and in WT primary hepatocytes after siNC or si*Cad*-mediated knockdown (**B**). The hepatocytes received a 24 hr exposure to glucose and palmitate as indicated before the 6 hr incubation in glucose production media that contained 2.5 mM glutamine, 5.0 mM lactate and 0.5 mM pyruvate (n = 3 per group). (**C**) Hepatic glucose production (HGP), gluconeogenesis (GNG), and glycogenolysis (GLY) in primary hepatocytes from CTL and KO mice. The hepatocytes received a 24 hr exposure to glucose and palmitate as indicated before the 6 hr incubation in glucose production media that contained 20 mM lactate and 2 mM pyruvate (n = 3 per group). (**D-F**) Hepatic glucose production (HGP), gluconeogenesis (GNG), and glycogenolysis (GLY) in primary hepatocytes from CTL and KO mice. The hepatocytes received a 24 hr exposure to glucose and palmitate as indicated before the 6 hr incubation in glucose production media that contained 20 mM lactate and 2 mM pyruvate. Half of the hepatocytes were supplemented with 10 µM uridine after cell attachment until the end of study (n = 3 per group). (**G-J**) Primary hepatocytes from CTL and KO mice were supplemented with or without 10 μM uridine for 24 hr and then incubated with glucose production media that contained 2.5 mM glutamine (100% [U-^13^C_5_]-glutamine), 5.0 mM lactate and 0.5 mM pyruvate for 6 h. Mass isotopomer analysis of glucose and uridine in cells (**G & I**) and media (**H & J**) (n = 3 per group). Basal media without uridine supplementation is glucose-free and uridine-free, while basal media with 10 μM uridine supplementation contained uridine at the level indicated by the dashed line in **J**. (**K**) Hierarchical clustering of metabolites in the cells from the study in (**G-J**) showing three 3 groups based on pattern (n = 3 per group). ^*^p < 0.05, ^**^p < 0.01, ^***^p< 0.001, ^****^p< 0.0001; ns, no significance. Error bars denote SEM. (**L**) A model of gluconeogenesis regulation by pyrimidine synthesis. In the liver, fasting suppresses pyrimidine synthesis to increase gluconeogenesis. Concomitant reduction in biliary excretion by fasting helps maintain blood uridine at an elevated level, which synergistically stimulates hepatic gluconeogenesis to prevent hypoglycemia. Obesity suppresses pyrimidine synthesis due to increased uridine output from adipocytes. As a consequence, gluconeogenesis is stimulated regardless of calorie intake or restriction, which then promotes insulin resistance and more obesity.

To further investigate the synergistical action of uridine and Cad loss on gluconeogenesis, [U-^13^C5] glutamine was used as a tracer to study uridine treatment effects on cellular metabolism in the control and Cad KO hepatocytes. After 6-hour culture in the media, the cellular glucose levels were increased by Cad KO, whereas, uridine combined with Cad KO showed the highest cellular glucose among all the groups (**Fig. 8G**). Media glucose contents, which revealed higher fractions of newly synthesized glucose including the M+3 isotopologue, recapitulated the profile of cellular glucose (**Fig. 8H**), confirming gluconeogenesis is increased most by Cad KO plus uridine. Both cellular and media uridine were reduced in Cad KO hepatocytes, confirming that *de novo* pyrimidine synthesis is suppressed by Cad KO (**Fig. 8I-J**). When uridine was not supplemented, the cellular uridine (M+0) in Cad KO was 27% of the control cells (**Fig. 8I**, CTL vs. KO), whereas when uridine was supplemented at 10 μM, the cellular uridine (M+0) in Cad KO was 44% of the control (**Fig. 8I**, CTL vs. KO Uridine 10 μM), suggesting partial restoration of cellular uridine in Cad KO hepatocytes by exogenous uridine. In control cells, the same concentration of uridine led to a 28% increase in cellular uridine (**Fig. 8I**, CTL vs. CTL Uridine 10 μM), suggesting that the control hepatocytes also can increase its cellular uridine concentration when exogenous uridine is supplied. Because the basal media is uridine-free, detection of uridine in conditioned media from the control and KO cells indicates hepatocytes actively release uridine (**Fig. 8J**). When cultured without uridine supplementation, control hepatocytes had media uridine up to 36% of the uridine in basal media, whereas the Cad KO hepatocytes only had media uridine up to 3% of the uridine in basal media (**Fig. 8J**), suggesting that *de novo* pyrimidine synthesis in WT hepatocytes was responsible for the uridine released in the culture media. When cultured with 10 μM uridine, control hepatocytes accrued almost 2-fold more uridine in the media compared to WT hepatocytes without uridine supplementation (**Fig. 8J**), confirming that uridine supplementation at 10μM did not cause a supraphysiological high level of uridine. As expected, Cad KO cells actively took up uridine when uridine was supplemented, so the uridine in culture media from Cad KO hepatocytes was only 14% of the basal media (**Fig. 8J**). In summary, uridine supplementation increases uridine uptake in both WT and Cad KO hepatocytes.

Gluconeogenesis relies on tricarboxylic acid cycle for substrate shuffling and ATP production and in this is a process opposite to glycolysis. We examined the metabolites involved in glucose metabolism and tricarboxylic acid cycle for differential enrichment between control and Cad KO cells (**Fig. 8K**). In Cad KO cells, pyruvate and lactate levels were lower than control cells, suggesting that the consumption of pyruvate and lactate is increased in Cad KO hepatocytes as substrates for gluconeogenesis. In contrast to pyruvate and lactate, metabolites including 2,3-diphosphoglycerate and ⍺-KG were increased in Cad KO cells but not by uridine alone in WT cells, while others including fructose-6-phosphate and citrate showed an increase by uridine but not by Cad KO alone. Notably, the overall profile showed that uridine supplementation to Cad KO hepatocytes resulted in more metabolites to be elevated than Cad KO or uridine treatment alone. This supports the view that the synergy of uridine and Cad loss in promoting gluconeogenesis occurs at the substrate level of carbohydrate metabolism and in a cell autonomous manner. Consistent with the metabolomic data, the pathways enriched of those metabolites, that is, glycolysis, TCA cycle, and pentose phosphate pathway were upregulated in the livers from Cad KO mice in fed state (**Fig. S8A-C**). Thus, the stimulation of gluconeogenesis by Cad KO is associated with increased liver capacity for carbohydrate metabolism.

## Discussion

### Hepatic *de novo* uridine biosynthesis is dispensable for maintenance of circulating uridine but indispensable for regulation of bile uridine

Most tissues and cells rely on the blood for their uridine supply [40]. The liver is considered the primary organ responsible for regulating plasma uridine levels, overseeing the *de novo* biosynthesis within hepatocytes and the clearance of uridine by Kupffer cells[17, 41]. Our study indicates that hepatocyte Cad KO, which efficiently reduced hepatic uridine release (**Fig. 7A**), did not cause a decrease in blood uridine (**Fig. 7E**). Thus, suppression of *de novo* pyrimidine synthesis in the liver does not result in a decrease in circulating uridine. Moreover, HFD caused an increase in blood uridine in the Cad KO mice and the control mice (**Fig. 7E**), indicating that the uridine supply from the liver is not relevant in relation to diet-induced elevation in blood uridine. This finding is consistent with our recent report that adipocytes are capable of *de novo* uridine synthesis and serve as a key source for uridine supply in fasted mice [14]. Indeed, the hepatic Cad KO mice had two times the level of adipocyte Cad (**Fig. 7J-K**), indicating that the adipocyte capacity of uridine supply is increased so that the circulating uridine levels are maintained unchanged in the hepatocyte-specific Cad KO mice.

The liver uridine content in the Cad KO mice, however, was lower than control group when fed on HFD but not on NCD (**Fig. 7D**), suggesting a diet-dependent response. In contrast, bile uridine concentration in male Cad KO mice was higher than control group when fed on HFD but not on NCD (**Fig. 7F**). The reciprocal changes in liver uridine and bile uridine by Cad KO suggest uridine enrichment in bile is enhanced when *de novo* uridine synthesis is suppressed. Since uridine facilitates phospholipid synthesis in enterocytes [42, 43], the increase of bile uridine by HFD in the Cad KO mice might increase intestine lipid absorption. Concurrent with this hypothesis, Cad KO mice gain weight faster than control animals on HFD but not on NCD (**Fig. 2A & Supplemental Fig.2A**). The higher weight gain in Cad KO mice was not tied to increased food intake (**Fig. 5E**). Notably, the increase in bile uridine was concomitant with an increase of 12⍺-hydroxylated bile acid relative to 12 non-hydroxylated bile acid, which is known to increase the efficacy of lipid uptake [38]. The combined increase in bile uridine and shift in bile acid composition might explain why Cad KO-facilitated weight gain was significant only in HFD group and not in NCD mice.

### Suppression of *de novo* hepatic pyrimidine synthesis promotes diet-induced obesity and insulin resistance

We found that both gene expression and protein levels of Cad were reduced in the liver from HFD mice, suggesting diet-induced obesity is associated with a reduction in liver uridine biosynthesis. High glucose and high palmitate, which mimics the metabolic changes and stress in obesity, caused a reduction in Cad in primary hepatocytes, indicating that obesity suppresses Cad expression in hepatocytes. To examine the metabolic consequence of Cad downregulation, we generated a mouse model of hepatocyte-specific Cad KO. In addition to higher weight gain and fat mass, liver TG and cholesterol levels were also increased in the Cad KO mice on HFD, a metabolic derangement similar to metabolic syndrome in humans. Clamp experiments confirmed that the Cad KO mice were more insulin resistant than control mice on HFD, indicating that a reduction in hepatic uridine biosynthesis is a cause of insulin resistance in diet-induced obesity. Interestingly, hepatocyte-specific overexpression of UPP1 lowered liver uridine content and caused lipid accumulation [3]. Since low uridine and high lipid content were found in the livers from Cad KO mice (**Fig. 7D, 2E-F**), it appears that the liver uridine content, rather than blood uridine levels, is tied to hepatic lipid accumulation.

Using indirect metabolic calorimetry metabolic cages, we found that food intake was not significantly increased in Cad KO mice compared to the control group, suggesting that the higher rate of weight gain is not likely from increased food intake. This result is consistent with ours and others’ model in which elevated blood uridine acts as a signal of hunger [15, 18], whereas Cad KO mice do not show a difference in circulating uridine compared to the HFD-fed control mice. Respiratory exchange rates were decreased in the Cad KO mice, hinting at an increase in fatty acid oxidation. Despite increased fatty acid oxidation, the Cad KO mice still gain weight faster than the control mice on HFD, suggesting that their lipid absorption is increased, which is supported by the changes found in bile uridine levels and bile acid composition in the Cad KO mice (**Fig. 7F-H**).

### Hepatic uridine biosynthesis is coupled to gluconeogenesis and first-passage insulin clearance

*De novo* pyrimidine synthesis consumes ribose-5-phosphate and aspartate. This biosynthesis process is unfavored when glucose production is demanded in hepatocytes. Indeed, the *de novo* pyrimidine synthesis pathway was downregulated in the liver by overnight fasting [14]. We showed that Cad protein, the first and rate-limiting enzyme in the biosynthesis pathway, tracked with its transcript level and decreased 20% by fasting (**Fig. 1F**), supporting the hypothesis that uridine biosynthesis must be suppressed for hepatocytes to increase gluconeogenesis. In lieu of this, *de novo* pyrimidine synthesis could prevent excess gluconeogenesis in the fed state. This provides an explanation for why the fasting-refeeding regulation of pyrimidine synthesis occurs in hepatocytes and not in adipocytes. Since HFD induces a reduction in hepatocyte Cad with feeding, this implies that HFD will drive gluconeogenesis as a result of reduced pyrimidine synthesis.

*In vivo* examination of the impact of pyrimidine synthesis on gluconeogenesis revealed that blood glucose in Cad KO mice rose higher than the control group in response to CL316,243-induced lipolysis or exogenous gluconeogenic pyruvate (**Fig. 6C-F**). Key players in gluconeogenesis including Pgc1⍺, FoxO1, PCX, and PEPCK, were upregulated in the livers from Cad KO mice (**Fig. 6G-I**). *In vitro* experiments in primary hepatocytes indicated that Cad suppression increased gluconeogenesis under conditions mimicking metabolic stress in obesity (**Fig. 6M**). Therefore, our *in vitro* studies agree with the *in vivo* data, indicating that suppression of pyrimidine synthesis stimulates gluconeogenesis. Unexpectedly, uridine is found to potently increase gluconeogenesis in Cad KO hepatocytes (**Fig. 8D-F**), suggesting a synergy of uridine and Cad loss, which is further validated by the metabolomic profiling data (**Fig. 8G-K**).

Gluconeogenesis *in vivo* is regulated by insulin. Interestingly, our results suggest that hepatic Cad KO leads to a decrease in insulin clearance in the liver (**Fig. 4A-F**). β-cell secreted insulin is first cleared up to 50% in the liver before it enters the circulation. Lowering insulin clearance at the liver leads to systemic hyperinsulinemia which, in turn, promotes obesity and insulin resistance. Therefore, reduced clearance of hepatic insulin has been proposed as a risk factor for diabetes [44]. Our findings indicate that *de novo* pyrimidine synthesis is involved in the control of insulin degradation by CEACAM1(**Fig. 4G-H**). Thus, our work uncovers a significant role for hepatocyte *de novo* pyrimidine synthesis in both gluconeogenesis and insulin clearance.

### Remodeling of uridine homeostasis by obesity promotes obesity and T2D

The contribution of obesity to insulin resistance extends beyond the well-characterized disruption of insulin signaling. It encompasses complex interactions among multiple metabolic circuits, key nutrients, and bioactive metabolites [45]. The results herein reveal how uridine, the most abundant nucleoside in circulation, contributes to the etiology of obesity and T2D. We demonstrated that uridine homeostasis, from circulating level to biosynthesis capacity, is profoundly altered in diet-induced obesity. The liver, but not adipose tissue, is suppressed in *de novo* pyrimidine biosynthesis by HFD. From studying the metabolic sequalae of hepatic CAD KO, we found that suppression of *de novo* pyrimidine synthesis in hepatocyte leads to increased gluconeogenesis. The Cad KO hepatocytes still release uridine when cultured, suggesting that a uridine reservoir might exist in hepatocytes. Indeed, cellular RNA is reported as a source of uridine [46]. Notably, high glucose and high palmitate, as a mimic of metabolic stress in obesity, reduced uridine release from control hepatocytes but not the Cad KO ones, indicating the Cad-mediated uridine synthesis, but not the hepatic uridine reservoir, is sensitive to the metabolic stress.

Unexpectedly, uridine synergized with Cad deletion, indicating that elevated uridine will further increase hepatocyte gluconeogenesis when pyrimidine synthesis is suppressed, a setting akin to that observed in obesity and fasting. The pro-gluconeogenic effect of uridine is consistent with findings that uridine depletion reduced mitochondrial pyruvate oxidation [47], suggesting that uridine-facilitated mitochondrial ATP production is a potential mechanism for its synergistical action with Cad deletion. Together, the findings indicate that suppression of *de novo* pyrimidine synthesis in hepatocytes is a key event of uridine homeostasis remodeling which links uridine metabolism to gluconeogenesis (**Fig. 8L**). The reciprocal change in pyrimidine synthesis and gluconeogenesis during fasting suggests that uridine homeostasis is integral to fasting adaptation and is beneficial for survival under starvation. In contrast, diet-induced obesity increased uridine production and release by adipocytes uridine, which downregulated hepatocyte Cad to suppress pyrimidine synthesis. As a result, concomitant elevation of blood uridine and suppression of hepatocyte pyrimidine synthesis is induced in obesity, which synergistically stimulates gluconeogenesis, aggravating the development of obesity and T2D.

The circulating uridine is typically maintained within a narrow range between 3–8 μM in mammals [11, 48]. Because high level of uridine triggers misincorporation of uridine into DNA, high rates of spontaneous tumorigenesis are observed in Upp1 knockout mice that show also elevated blood uridine [13]. Thus, theologically, maintenance of blood uridine within a narrow range likely protects from high-rate tumorigenesis. Since obesity leads to an increase in adipocyte uridine supply, suppressing uridine synthesis in hepatocytes becomes a critical response to prevent spontaneous tumorigenesis. Yet, hepatocyte pyrimidine synthesis suppression, promotes gluconeogenesis and ultimately triggers obesity and T2D. Therefore, our findings suggest a new paradigm for the etiology of metabolic deterioration in diet-induced obesity, in which uridine homeostasis remodeling promotes the progression of obesity and T2D.

### Potential problems and future directions

Both adipocytes and hepatocytes are capable of *de novo* pyrimidine synthesis, but only hepatocytes are suppressed by HFD for pyrimidine synthesis. To uncover the underlying mechanism will help identify new strategies to prevent the suppression of uridine synthesis in hepatocytes and, therefore, to prevent excessive gluconeogenesis. In addition to adipocytes, macrophages contribute to local uridine supply [49]. Diet-induced obesity is associated with chronic inflammation which might induce an elevation in liver uridine from macrophage-mediated local uridine release. It remains elusive how chronic inflammation affects uridine homeostasis and hepatic uridine synthesis.

Food intake lowers blood uridine, which might serve as a mechanism for the reactivation of *de novo* pyrimidine synthesis in hepatocytes upon refeeding. However, unlike the other cell types, hepatocytes receive portal blood which contains uridine absorbed from intestine. Thus, the uridine concentration in the portal vein will affect the uridine content in the liver and, ultimately, gluconeogenesis. Meanwhile, cholestasis and liver steatosis might affect hepatocyte release of uridine through the bile, which in turn feedback to suppress hepatic pyrimidine synthesis. Studying the effect of obesity on the dynamic changes in portal uridine will completement the studies described herein and validate our new paradigm about the role of uridine homeostasis in obesity and T2D development.

The phenomenon where impaired insulin signaling blocks one of its actions (decrease of gluconeogenesis) while promoting another action (lipogenesis) is termed selective insulin resistance, and is displayed in the insulin-resistant state in rodent models of T2D [50]. A bifurcation point in the insulin signaling in the liver is suggested after Akt and prior to the mTORC1 complex [51]. The hepatocyte Cad KO mice fully recapitulate this selective insulin resistance, indicating that suppression of *de novo* pyrimidine synthesis is a cause for the selective insulin resistance. It remains to be seen how insulin-mediated suppression of gluconeogenesis is selectively blocked by loss of CAD while the insulin signaling upstream of the bifurcation point remains unaffected.

We used HFD feeding in mice to model the development of obesity and prediabetes. Whether the uridine homeostasis remodeling found in these experiments is significant for individuals with established T2D should be pursued. Since gluconeogenesis is a primary driver of hepatic glucose production in patients with T2D [52] and increased uridine is found in clinical T2D [9], determining the regulation of hepatocyte pyrimidine synthesis and the disruption of uridine homeostasis in T2D will contribute to our understanding of uridine in the pathophysiology of T2D.

## Materials and methods

### Mice and tolerance tests

C57BL/6N mice were housed in individually ventilated cages with constant temperature on a 12 hr dark/light cycle with lights on at 6 AM to 6 PM and free access to water and food. Mice were maintained on a rodent chow diet (LabDiet #5053) for normal chow diet (NCR) to 12 weeks old and were switched to a high-fat diet (Research Diet, D12492) for HFD-induced obesity. The Cad^f/f^ mice were derived from an ES clone from UCDAVIS KOMP Repository at the transgenic core in the University of Texas Southwestern Medical Center [18]. Hepatocyte-specific Cad KO mice were generated by breeding albumin-Cre transgenic mice (Jackson Laboratory) to Cad^f/f^ mice. Sex and age-matched littermates of hepatocyte Cad KO mice that were homozygous for Cad^f/f^ but lacking the albumin-Cre transgene were used as controls. Unless specified, the data were collected from male mice if not specified. Animal care and experimental protocols were approved by the Institutional Animal Care and Use Committee of City of Hope (Duarte, CA).

Glucose tolerance tests were conducted in mice after 4∼6 hr fasting as described [53]. Glucose (Sigma) was administrated by gavage (25% solution) at 2.5 g/kg bodyweight. For pyruvate tolerance tests, mice received 2 g/kg body weight of sodium pyruvate through intraperitoneal injection after fasting for 16 hours. For stimulated lipolysis, mice were injected subcutaneously with 1 mg/kg body weight CL316,243 (Sigma) and food was withdrawn after injection. For all the tests, blood was collected via tail nick using heparinized capillary tubes (Fisher) at time points of 0, 15, 30, 60, and 120 minutes after treatment. Plasma samples were centrifuged and stored at -20°C.

### Body composition, indirect calorimetry, and hyperinsulinemic-euglycemic clamp

All the studies were conducted by the Comprehensive Metabolic Phenotyping Core at City of Hope. Body composition was precisely measured using quantitative magnetic resonance technology (EchoMRI 3-in-1, Echo Medical Systems). Indirect calorimetry was conducted in metabolic cages (Phenomaster, TSE Systems). Mice were individually housed with *ad libitum* access to food and water throughout the experiment. The mice were allowed to acclimatize to the metabolic cages for 1 day prior to data collection. Data was analyzed using CalR software (https://CalRapp.org/). Hyperinsulinemic-euglycemic clamp was performed as described [54–56]. Briefly, carotid artery and jugular vein catheters were surgically placed for sampling and infusions 5 days before the study. Insulin clamps were performed on mice fasted for 5 h. Baseline blood or plasma parameters were obtained in blood samples collected at −15 and −5 minutes. Insulin was infused (4 mU/kg/min) at time 0 and continued for 155 minutes. Arterial glucose levels were monitored every 10 minutes to adjust the glucose infusion rate. [3-^3^H] glucose kinetics were determined at -15 minutes, 15 mininutes and from 80 to 120 minutes. [2-^14^C] deoxyglucose (13 μCi) was administered at 120 minutes. Then blood was taken at 2, 5, 15, 25 and 35 minutes for the determination of [2-^14^C] deoxyglucose. At the end of clamp, mice were anaesthetized and tissues were collected.

### Histology and metabolite measurements

Mouse livers were harvested and fixed in 10% formalin for 40 hr and further processed for hematoxylin and eosin (H&E) staining, trichrome staining, Periodic acid–Schiff (PAS) staining, and oil red O (ORO) staining by the Research Pathology Core at City of Hope. Stained sections were imaged using a BZ-X800 (Keyence) microscope and captured using 20x objective lens. For immunofluorescence, the paraffin sections of pancreas were de-paraffinized and subjected to antigen retrieval (Vector) and blocking in PBS with 2.5% BSA and incubated with primary antibodies at 4°C overnight. Primary antibodies used were guinea pig anti-insulin (Dako, A0564) and mouse anti-glucagon (Abcam, ab10988). Secondary antibodies used were Alexa Flour 568-labeled (Invitrogen, A11031) and Alexa Flour 488-labeled (Invitrogen, A11073). Slides were imaged using a BZ-X800 microscope after mounting with ProLong™ Gold Antifade Mountant with DAPI (Invitrogen, P36931) and captured using a 20x objective lens.

Blood glucose was measured using a glucometer (Ascensia Contour next). The HOMR-IR index was calculated as FBG✕FINS/22.5, where FBG is the fasting blood glucose level (mmol/L), and FINS is the fasting insulin level (mU/mL). Serum cholesterol and triglyceride levels were detected using Infinity Cholesterol (Thermo Scientific) and Infinity Triglyceride (Thermo Scientific) kits. Serum NEFA and glycerol levels were measured using a NEFA kit (FujiFilm-Wako) and free glycerol reagent (Sigma). Plasma ALT and AST levels were measured using a liquid ALT reagent set (Pointe) and AST colorimetric activity assay kit (Cayman). Plasma insulin and C-peptide levels were measured using an ultra-sensitive mouse insulin ELISA kit (Crystal Chem) and mouse C-peptide ELISA kit (Crystal Chem). Frozen liver tissues (40 mg) were used for lipid extraction and measurement as described previously [57]. Frozen liver tissues (20mg) were used for liver glycogen measurement as described [58]. Bile samples (2 ul) were analyzed for 30 bile acids as described [59].

### Uridine measurement by mass spectrometry

Uridine contents in frozen liver tissues were determined at the Integrated Mass Spectrometry Core (IMS) at the City of Hope Comprehensive Cancer Center. Metabolite extraction was performed by adding extraction buffer (acetonitrile, methanol, 3:1, v/v, 10 ul of ^13^C5 uridine as internal standard) to the tissue and homogenized in a Precellys homogenizer with a cryolys unit. Homogenized samples were clarified by centrifugation at 13000 rpm for 15 minutes at 4°C. The clear supernatant was diluted with equal amounts of water and 3 µL was used for LC-MS/MS analysis. Analytes were separated on Thermo Vanquish UPLC as described [60, 61] and analyzed on a Thermo TSQ-Altis mass spectrometer.

Plasma, bile, and cell culture medium uridine concentrations were determined by targeted liquid chromatography mass spectrometry (LC/MS).10 µM of Uridine·H_2_O, ^13^C9, ^15^N2, 96-98% (Cambridge Isotope Laboratories) solution in PBS was added to samples and incubated at room temperature for 1 hr. 450 µL of 80% methanol was added and clarified by centrifugation at 14,000 × g for 30 minutes at 4°C. 400 µL of the methanolic supernatant was dried on a frozen speed vac. The dried samples were redissolved in 0.1% formic acid in water inside of 0.2-µm PTFE filter vials (Thomson Carlsbad). LC/MS analysis was conducted on an Orbitrap Exploris 480 mass spectrometer equipped with an Atmospheric Pressure Chemical Ionization (APCI) source and a Vanquish UPLC system (ThermoFisher Scientific). Uridine was eluted from a Cadenza CD-C18 column (100×2 mm, 3 µm, Imtakt USA) by isocratic flow of 0.1% FA at 0.4 µL/min. Parallel Reaction Monitoring LC/MS in negative APCI mode selected and fragmented the parent ions of uridine and SIS at m/z 243.0623 and 254.0865, respectively. The resulting fragment ion chromatograms for samples and standard curves were analyzed with Skyline v23.1 [62].

### Untargeted metabolomic profiling

Untargeted metabolomics for liver tissue was conducted by the IMS at City of Hope’s Comprehensive Cancer Center using a previous method [63]. Briefly, 50 ± 1 mg liver tissues were homogenized, and metabolites were extracted using acetonitrile: methanol: water (11:33:20, v/v/v) spiked with isotopically labelled internal standards (ISTD, ^13^C Phenyl alanine, D4 Succinic acid, D8 Valine, ^13^C6 Adipic acid). Metabolite extracts were clarified and used for LC-MS/MS analysis. System suitability was determined using a plasma metabolite extract. Pooled QC was used for retention time drift correction and determining analytical variation during data acquisition and correction. Metabolites were subjected to RP and HILIC chromatographic separation using a Dionex 3000 UPLC and MS analysis using an Orbitrap Fusion Lumos mass spectrometer (Thermo) in positive and negative modes.

Untargeted metabolomics for mouse plasma was conducted by MD Anderson Cancer Center Metabolomics Core. Metabolites were extracted using ice-cold 0.1% Ammonium hydroxide in 80/20 (v/v) methanol/water. Extracts were centrifuged at 17,000 g for 5 minites at 4°C. Supernatants were dried by evaporation under nitrogen. Dried extracts were dissolved in deionized water, and injected for analysis by ion chromatography (IC)-MS. IC mobile phase A (MPA; weak) was water, and mobile phase B (MPB; strong) was water containing 100 mM KOH. A Thermo Scientific Dionex ICS-5000+ system included a Thermo IonPac AS11 column (4 µm particle size, 250 x 2 mm) with column compartment kept at 30°C. The autosampler tray was chilled to 4°C. The mobile phase flow rate was 350 µL/min and a gradient from 1mM to 100mM KOH was used. Data were acquired using a Thermo Orbitrap Fusion Tribrid Mass Spectrometer under ESI negative ionization mode at a resolution of 240,000.

### Primary hepatocytes isolation, treatment, siRNA for Cad knockdown, and glucose production assay with or without tracer

Primary hepatocytes were isolated from 8- to 12-week-old male mice as described [64, 65]. Cell viability was determined using a cell counting kit-8 (CCK8, Sigma) and 4 x 10^5^ cells per sample were counted. For uridine treatment, hepatocytes post attachment were cultured with 10 µM uridine (Sigma) until the cells were harvested at the end of study. For PALA treatment, hepatocytes post attachment were cultured with PALA [14] at concentrations of 0, 0.1, 10, and 100 µM. For knockdown studies, primary hepatocytes were transfected with siRNAs using siLentFect™ Lipid Reagent (Bio-Rad). For glucose production assay, primary hepatocytes were washed sequentially with prewarmed PBS and serum-free DMEM media before cultured in serum-free DMEM with specified glucose (1.0 g/L or 4.5g/L) ± palmitate (180 µM) for 24 hr. The hepatocytes were washed twice with prewarmed PBS and cultured in glucose production media (glucose-free DMEM without phenol red, 100-unit Penicillin and 0.1 mg/ml streptomycin) ± the indicated gluconeogenic substrates for 6 hr. The glucose in the culture medium was determined using a colorimetric assay (Sigma) and normalized to the cellular protein concentration measured by BCA assay (Pierce). Total hepatic glucose production was calculated from cells cultured with gluconeogenetic substrate, while glycogenolysis was calculated from cells cultured without gluconeogenetic substrate, and gluconeogenesis was calculated as the difference between the total hepatic glucose production and glycogenolysis [66]. For the tracer study, [U-^13^C5]-glutamine (2.5 mM, Cambridge Isotope Laboratories) was supplemented to glucose production assay for 6 hr. The cell lysate and culture media were subjected to targeted analysis of polar metabolites by high-resolution mass spectrometry/ion chromatography (HRLC/IC-MS) at MD Anderson Cancer Center Metabolomics Core.

### Western Blotting

Protein lysates were prepared in RIPA lysis buffer supplemented with protease and phosphatase inhibitor (Thermo Scientific). Lysate protein concentrations were determined by the BCA assay (Thermo Scientific). 20 μg of protein was resolved on 4-20% Tris-glycine gels (Bio-Rad), and transferred to nitrocellulose membranes (LI-COR). Protein expression was confirmed by anti-GAPDH (Fitzgerald, G109a), anti-CAD (Cell Signaling Technology, 11933), anti-CEACAM1 (Cell Signaling Technology, 14771), and anti-PCK1 (Cell Signaling Technology, 12940). Secondary antibodies used were IRDye 800-conjugated anti-rabbit (Li-Cor, 925-32211), Alexa Fluor 800-conjugated anti-mouse (Invitrogen, A32730), and IRDye 680-conjugated anti-mouse secondary antibodies (Li-Cor, 926-68070).

### RNA isolation for quantitative PCR and bulk RNA sequencing

Total RNAs from 50 mg frozen mouse tissues and primary hepatocytes were isolated with an Aurum Total RNA Fatty and Fibrous Tissue kit (Bio-Rad) and a Quick-RNA Microprep kit (Zymo Research). For each sample RNA (250ng) was used for reverse transcription (PrimeScript, Takara). cDNA samples were then diluted tenfold with ddH_2_O for quantitative PCR using iTaq Universal SYBR Green Supermix (Bio-Rad) and values were normalized with the 18s rRNA using ΔΔ–Ct method. Primer sequences are listed in supplementary Table 1 (Table S1). Total RNA from liver tissues was used for RNA sequencing (RNA-seq) analysis (Admera health).

### Statistical analyses

The nonparametric, two-tailed Student’s t test was used for two-group analyses. One-way or two-way analyses of variance (ANOVA) were used for multiple group analysis. Statistical analysis was conducted between genotypes unless specified. Data were reported as the mean±SEM. A p value < 0.05 was considered significant. All analyses were performed using GraphPad Prism Software 10.1.0 (GraphPad Software, Inc.).

## Supplementary materials

Supplementary table 1 and Supplementary figures 1-8.

## Acknowledgements

We thank Dr. Charles Brenner and Dr. Sarah Shuck for consultation for mass spectrometry analysis, Dr. Nisha Sharma for help on bile acids sample preparation, and Dr. Jeffrey Isenberg for critical reviewing of the manuscript. We thank the staff of the Comprehensive Metabolic Phenotype Core for their contributions to this work. We acknowledge the support of the Integrated Mass Spectrometry Core at City of Hope Comprehensive Cancer Center which is supported by the National Cancer Institute of the National Institutes of Health under award number P30CA33572.

## Author contributions

L.F. conceived of and conducted experiments with help from P.Z., I.N-S., Y.W., C.J., K.W., Y.P., and L.D. Uridine measurements were conducted by P.H.P. and M.K. RNA sequencing and public data analysis was conducted by Q.X. The clamp and metabolic cage experiments were assisted by P.F. Bile acids measurement was assisted by L.J. and W.H.. Metabolomic profiling analysis and tracer studies were assisted by P.L.L. and L.T.. Mouse model characterization was assisted by P.E.S. and Z.V.W.. Y.D. conceived of and designed the study, and wrote the manuscript. All the authors commented on and approved the manuscript.

## Data availability

All data related to this work can be found in the paper or supplemental materials.

