## Supplemental data for "*De Novo* Hepatic Pyrimidine Synthesis Regulates Systemic Energy Homeostasis"

**Supplementary Table 1. Primer sequences**

| Gene | Forward (5'-3') | Reverse (5'-3') |
| --- | --- | --- |
| *Cad* | CTGCCCGGATTGATTGATGTC | CTGCCCGGATTGATTGATGTC |
| *Dhodh* | TCTTCACCTCTTACCTGACAGC | CATGTTGGAGTCCTGAAACGTA |
| *Umps* | TGGCTGAGGAGCACTGTGAA | CGGTTAGCCGCTGCAAGTAT |
| *Upp1* | CCAACATCTGTGCAGGCACT | AACTCCGGCTTGAAGCACTC |
| *Upp2* | AGAGCTGGTGTCAGGAACAT | AGATTAGAAGCTGTGGCCGT |
| *Pgc1α* | AAGCGAAGAGCATTTGTCA | GAGGGTCATCGTTTGTGGT |
| *Foxo1* | GCAGCCAGGCATCTCATAA | CCTACCATAGCCATTGCAGC |
| *Hnf4α* | GCATGGATATGGCCGACTAC | TGTGGTTCTTCCTCACGCTC |
| *Pck1* | TGACAACTGTTGGCTGGCTC | GACATACATGGTGCGGCCTT |
| *Pcx* | GTGAGAT TGCCATCCGAGTG | TCTGCTCGCTCTGAGAGGAA |
| *G6pc* | GGCGCAGCAGGTGTATACTA | ATGCCTGACAAGACTCCAGC |
| *Pygl* | CCAGAGTGCTCTACCCCAAT | CAGCCACCACAAAGTACTCCT |
| *Lxrα* | CAGGAGTGTCGACTTCGCAA | GCCACCAGCTTCTCGATCAT |
| *Srebp1* | TGCCATTGAGAAGCGCTACC | TCCACTGCCACAAGCTGACA |
| *Srebp2* | AAGTCCTGCAGCCTCAAGTG | CCATCTGTCTTCAGCGTGGT |
| *Fasn* | AGGAGGTGGTGATAGCCGGT | GGTCCATTGTGTGTGCCTGC |
| *Scd1* | GCCGAGAAGCTGGTGATGTT | ATAGAGATGCGCGGCACTGT |
| *Dgat2* | GCCTGCAGTGTCATCCTCAT | TGGTGGTCAGCAGGTTGTGT |
| *Rn18s* | AGGGTTCGATTCCGGAGAGG | CAACTTTAATATACGCTATTGG |

### Supplementary Fig. 1

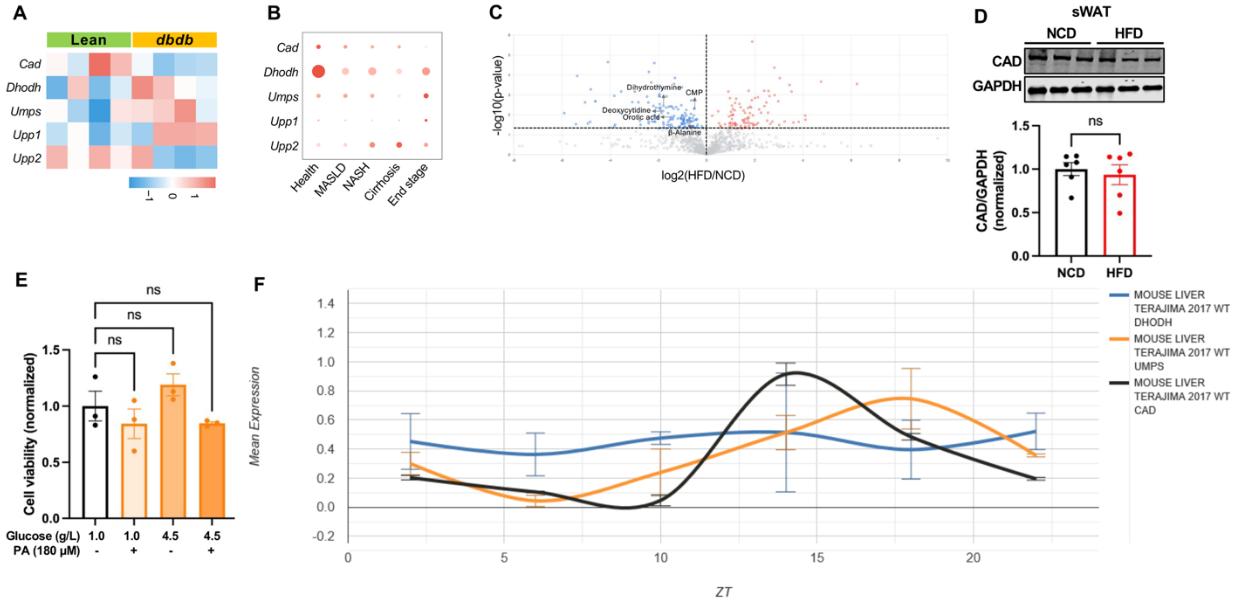


**Supplementary Figure 1. *De novo* hepatic pyrimidine biosynthesis is downregulated in T2D mice, in subjects with NASLD, by HFD feeding, and shows a circadian rhythm.**
(**A**) Heatmap of genes in pyrimidine metabolism generated with z-score from bulk RNA sequencing data. The raw data from the livers of *dbdb* mice and lean mice was retrieved from the dataset associated with study [20].
(**B**) Dotplot of genes in pyrimidine metabolism from hepatocytes. The raw data of single-nucleus RNA sequencing from liver biopsies was retrieved from the dataset associated with study [21].
(**C**) Volcano plot of metabolites in the livers from mice on NCD or after 15 weeks of HFD feeding (n = 3 per group). Tissues were harvested from random fed mice. Dotted lines denote the significance cut-off ($p < 0.05$).
(**D**) Western blot and quantification of Cad protein in the sWATs from mice on NCD or after 12 weeks of HFD feeding (n = 6 per group). Tissues were harvested from random fed mice.
(**E**) Cell viability of primary hepatocytes from WT C57BL/6 mice. The Hepatocytes were exposed to glucose and palmitate as indicated for 24 hr (n = 3 per group).
(F) Circadian rhythm profile of de novo pyrimidine biosynthesis genes in the livers from WT mice fed on chow. Data is retrieved from http://circadiomics.igb.uci.edu/ according to the study described [24]. ZT, zeitgeber time. ZT 0 is the "lights on" onset and ZT 12 is the "lights off" onset on a 12-hour light, 12-hour dark schedule. ns, no significance. Error bars denote SEM.

### Supplementary Fig. 2

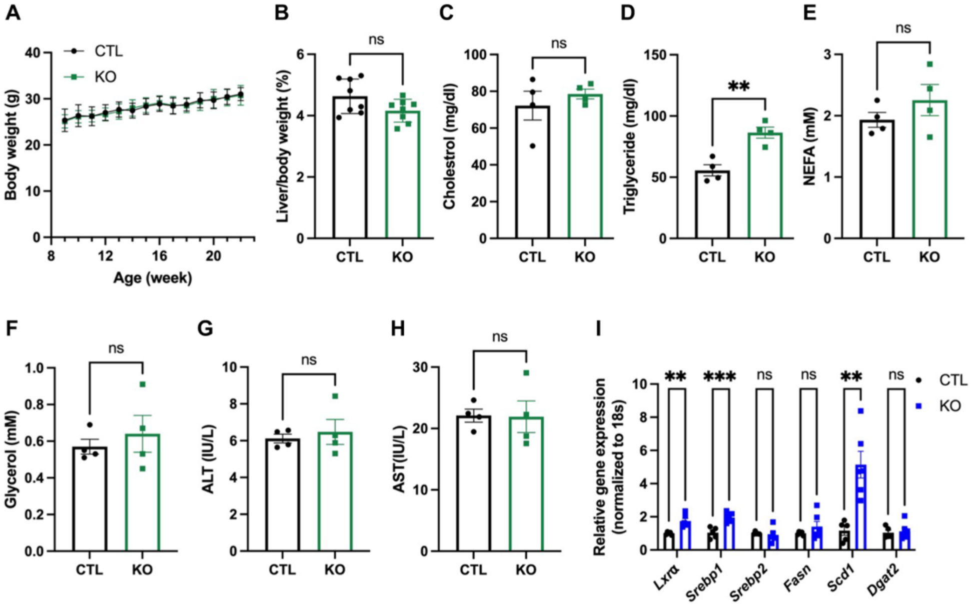


**Supplementary Figure 2. Hepatic Cad loss promotes weight gain in mice on HFD but not on NCD.**
(**A**) Weekly body weight for CTL (n = 9) and KO (n = 8) mice during 22 weeks of NCD feeding.
(**B**) Liver weight-to-body weight ratio in CTL and KO mice after 20 weeks of NCD feeding (n = 8 per group). Measurements were done after a 24h fast.
(**C-F**) Serum cholesterol (**C**), triglyceride (**D**), NEFA (**E**) and glycerol (**F**) in CTL and KO mice after 20 weeks of NCD feeding (n = 4 per group). Serum was collected after a 24 hr fast.
(**G-H**) Plasma ALT and AST in CTL and KO mice after 20 weeks of NCD feeding (n = 4 per group). Plasma was collected after a 24h fast.
(**I**) qPCR analysis of *de novo* lipogenesis pathway genes in the livers of CTL (n = 5) and KO (n =6) mice after 23 weeks of HFD feeding. Tissues were collected after a 24 hr fast. **$p < 0.01$, ***$p < 0.001$; ns, no significance. Error bars denote SEM.

### Supplementary Fig. 3

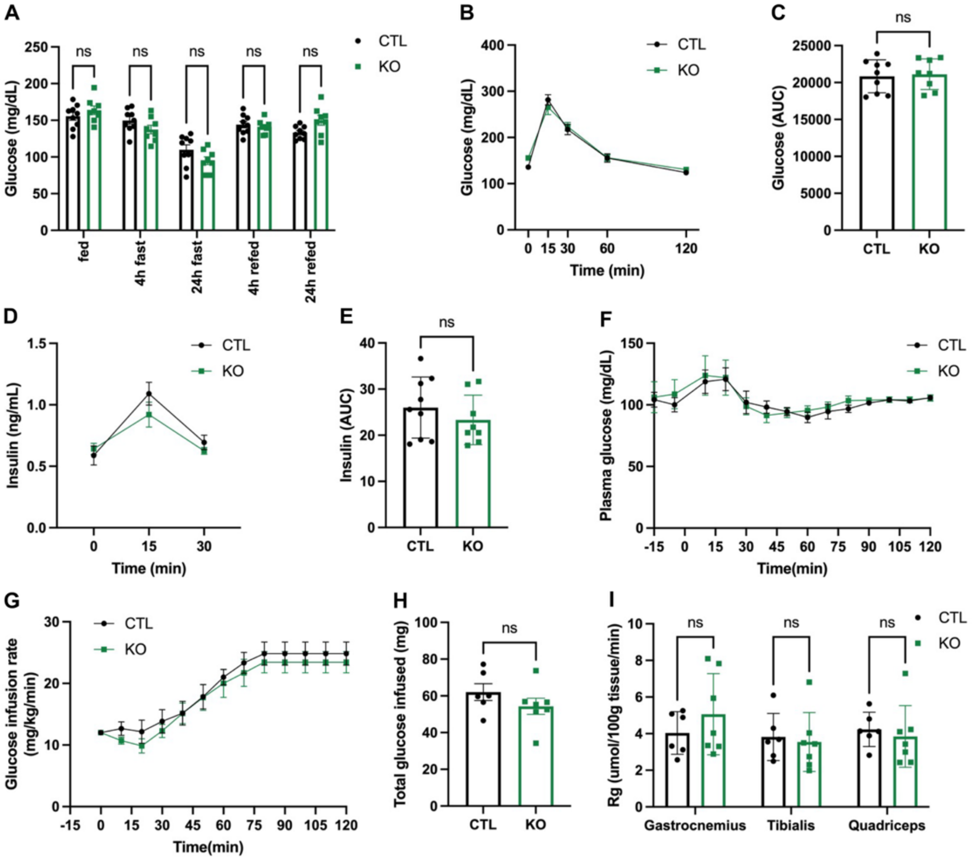


**Supplementary Figure 3. Hepatic Cad loss exacerbates insulin resistance in mice on HFD but not on NCD.**
(**A**) Blood glucose in CTL (n = 9) and KO (n = 8) mice during a fed/fast/refed study. The mice were fed on NCD for 12 weeks.
(**B-C**) Blood glucose response to glucose overload during an OGTT in CTL (n = 9) and KO (n = 8) mice after 10 weeks of NCD feeding.
(**D-E**) Blood insulin response to glucose overload in the OGTT in (**B**).
(**F**) Blood glucose levels at 15-minute intervals throughout hyperinsulinemic-euglycemic clamp for CTL (n = 6) and KO (n = 7) mice after 22 weeks of NCD feeding.
(**G-H**) Glucose infusion rate (**G**) and total glucose infused (**H**) during the hyperinsulinemic-euglycemic clamp study in (**F**).
(**I**) Gastrocnemius, tibialis anterior, and quadriceps muscle glucose uptake during the hyperinsulinemic-euglycemic clamp study in (**F**). ns, no significance. ns, no significance. Error bars denote SEM.

### Supplementary Fig. 4

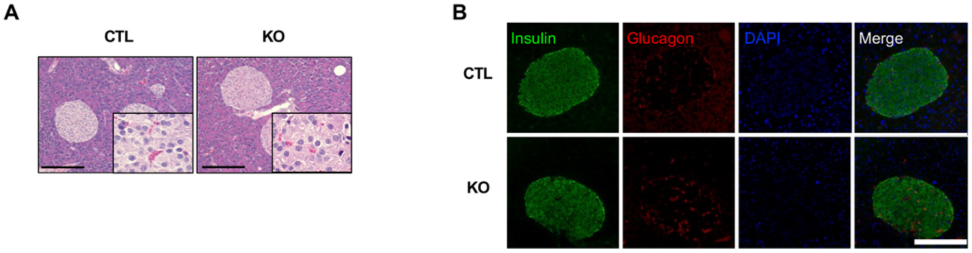


**Supplementary Figure 4. Hepatic Cad KO reduces insulin clearance in diet-induced obesity without affecting islets.**
(**A**) Representative hematoxylin-eosin staining of pancreata from CTL and KO mice after 20 weeks of HFD feeding. Scale bar, 200 μm. Insets are magnified 16 times.
(**B**) Representative images of immunofluorescent staining of insulin (green), glucagon (red), and DAPI (blue) of tissue sections from pancreata from CTL and KO mice after 20 weeks of HFD feeding. Scale bar, 200 μm.

### Supplementary Fig. 5

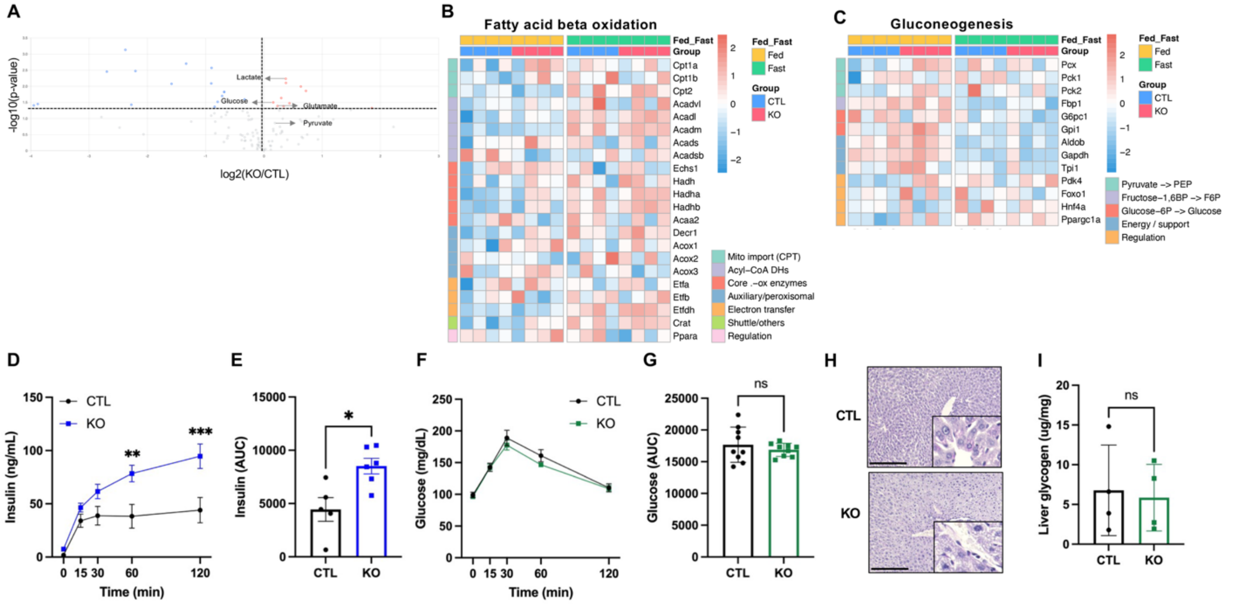


**Supplementary Figure 5. Impact of hepatic Cad KO on systemic energy homeostasis in HFD fed mice is not observed in NCD fed mice.**

(**A**) Volcano plot of blood metabolites from CTL and KO mice after 23 weeks of HFD feeding. Plasma was collected after a 24 hr fast and metabolites determined by LC-MS/MS (n = 3 per group). Dotted lines denote the significance cut-off ($p < 0.05$).

(**B-C**) Heatmap of genes in fatty acid beta oxidation (**B**) and gluconeogenesis (**C**) pathways. Heatmaps were generated with z-score from bulk RNA sequencing data. Liver tissues were collected from CTL and KO mice after 15 weeks of HFD feeding, fed or after a 24 hr fast (n = 4 per group).

(**D-E**) Blood insulin response to CL316,243 (**D**) and the AUC (**E**) in CTL (n = 5) and KO (n = 6) mice after 18 weeks of HFD feeding.

(**F-G**) Blood glucose response to pyruvate (**F**) and the AUC (**G**) in CTL (n = 9) and KO (n = 8) mice after 13 weeks of NCD feeding.

(**H**) Representative PAS staining of liver sections from CTL and KO mice after 20 weeks of NCD feeding. Scale bar, 200 μm. Insets are magnified 16 times. Tissues were collected after a 24 hr fast.

(**I**) Hepatic glycogen contents of CTL and KO mice after 13 weeks if NCD feeding (n = 4 per group). Tissues were collected after a 24 hr fast. ns, no significance. Error bars denote SEM. *$p < 0.05$, **$p < 0.01$, ***$p< 0.001$. Error bars denote SEM.

### Supplemental Fig. 6

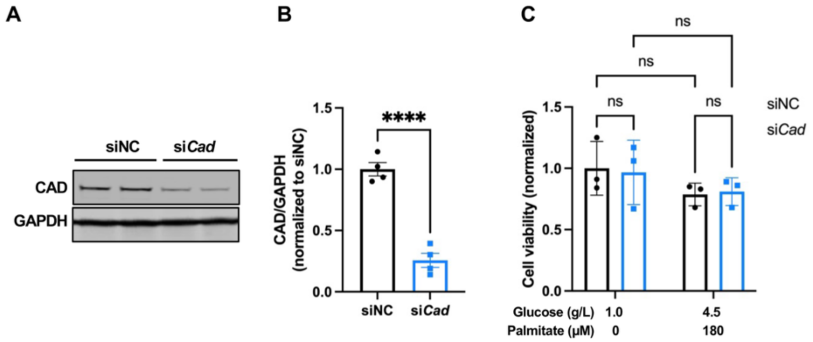


**Supplementary Figure 6. Hepatocyte Cad reduction by siRNA-mediated knockdown.**
(**A-B**) Western blot (**A**) and quantification (**B**) of Cad protein in WT primary hepatocytes after siNC or si*Cad*-mediated knockdown (n = 3 per group).
(**C**) Cell viability of WT primary hepatocytes after siNC or si*Cad*-mediated knockdown followed with a 24 hr exposure to glucose and palmitate as indicated (n = 3 per group). ****$p < 0.0001$; ns, no significance. Error bars denote SEM.

### Supplementary Fig. 7

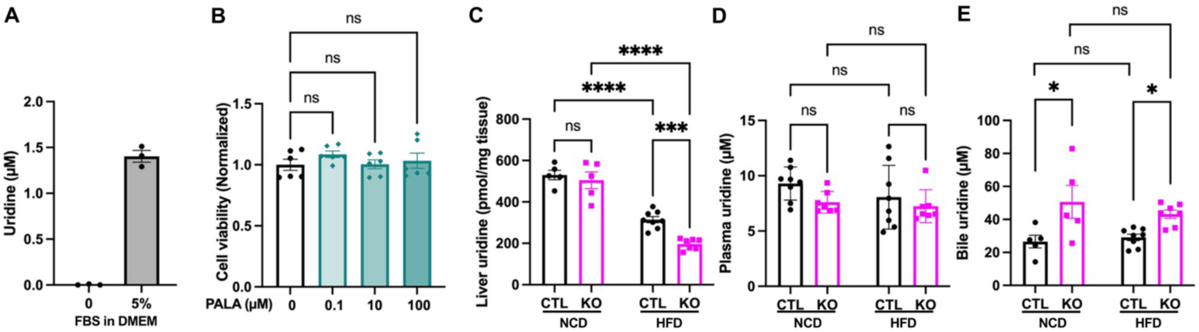


**Supplementary Figure 7. Impact of hepatic Cad KO on hepatic uridine production and circulating uridine levels in female mice.**

(**A**) Uridine levels in the DMEM with or without Fetal Bovine Serum (FBS) (n = 3 per group).

(**B**) Cell viability of hepatocytes after a 24 hr exposure to PALA at 0, 0.1, 10, 100 μM (n = 3 per group).

(**C**) Liver uridine contents of female CTL and KO mice on NCD or after 23 weeks of HFD feeding. Tissues were collected after a 24 hr fast (n = 5 ~ 11 per group).

(**D**) Plasma uridine levels of female CTL and KO mice on NCD or after 16 weeks of HFD feeding. Plasma was collected after a 24h fast (n = 7 or 8 per group).

(**E**) Biliary uridine levels of female CTL and KO mice on NCD or after 23 weeks of HFD feeding (n = 4 ~ 10 per group). *p < 0.05, ***p< 0.001, ****p< 0.0001; ns, no significance. Error bars denote SEM.

### Supplementary Fig. 8

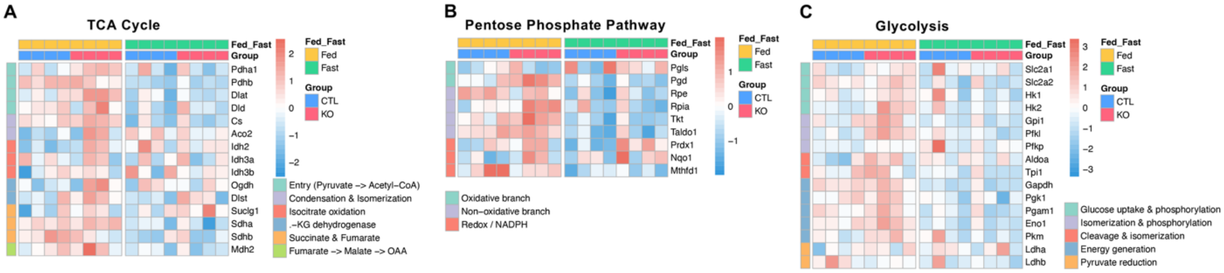


**Supplementary Figure 8. Uridine synergistically increase hepatic glucose production with hepatocyte Cad loss.**
(**A-C**) Heatmap of genes in TCA cycle (**A**), pentose phosphate (**B**), and glycolysis (**C**) pathways. Heatmaps were generated with z-score from bulk RNA sequencing data from the livers of CTL and KO mice after 15 weeks of HFD feeding. Liver tissues were collected from mice, fed or after a 24 hr fast (n=4 per group).
